# Contrasted reaction norms of wheat yield in pure vs mixed stands explained by tillering plasticities and shade avoidance

**DOI:** 10.1101/2022.10.27.514050

**Authors:** Meije Gawinowski, Jérôme Enjalbert, Paul-Henry Cournède, Timothée Flutre

**Affiliations:** Université Paris-Saclay, INRAE, CNRS, AgroParisTech, GQE - Le Moulon, 91190 Gif-sur-Yvette, France; Université Paris-Saclay, CentraleSupélec, MICS, 91190 Gif-sur-Yvette, France; Université Paris Cité, 75006 Paris, France

**Keywords:** plant-plant interactions, phenotypic plasticity, tillering, cultivar mixtures, bread wheat, yield components, shade avoidance syndrome

## Abstract

**Context:** Mixing cultivars is an agroecological practice of crop diversification, increasingly used for cereals. The yield of such cereal mixtures is higher on average than the mean yield of their components in pure stands, but with a large variance. The drivers of this variance are plant-plant interactions leading to different plant phenotypes in pure and mixed stands, i.e phenotypic plasticity.

**Objectives:** The objectives were (i) to quantify the magnitude of phenotypic plasticity for yield in pure versus mixed stands, (ii) to identify the yield components that contribute the most to yield plasticity, and (iii) to link such plasticities to differences in functional traits, i.e. plant height and flowering earliness.

**Methods:** A new experimental design based on a precision sowing allowed phenotyping each cultivar in mixture, at the level of individual plants, for above-ground traits throughout growth. Eight commercial cultivars of *Triticum aestivum* L. were grown in pure and mixed stands in field plots repeated for two years (2019-2020, 2020-2021) with contrasted climatic conditions and with nitrogen fertilization, fungicide and weed removal management strategies. Two quaternary mixtures were assembled with cultivars contrasted either for height or earliness.

**Results:** Compared to the average of cultivars in pure stands, the height mixture strongly underyielded over both years (-29%) while the earliness mixture overyielded the second year (+11%) and underyielded the first year (-8%). The second year, the magnitude of cultivar’s grain weight plasticity, measured as the difference between pure and mixed stands, was significantly and positively associated with their relative yield differences in pure stands (R^2^=0.51). When grain weight plasticity, measured as the log ratio of pure over mixed stands, was partitioned as the sum of plasticities in each yield component, its strongest contributor was the plasticity in spike number per plant (∼56% of the sum), driven by even stronger but opposed underlying plasticities in both tiller emission and regression. For both years, the plasticity in tiller emission was significantly, positively associated with the height differentials between cultivars in mixture (R^2^=0.43 in 2019-2020 and 0.17 in 2020-2021).

**Conclusions:** Plasticity in the early recognition of potential resource competitors is a major component of cultivar strategies in mixtures, as shown here for tillering dynamics. Our results also highlighted a link between plasticity in tiller emission and height differential in mixture. Both height and tillering dynamics displayed plasticities typical of the shade avoidance syndrome.

**Implications:** Both the new experimental design and decomposition of plasticities developed in this study open avenues to better study plant-plant interactions in agronomically-realistic conditions. This study also contributed a unique, plant-level data set allowing the calibration of process-based plant models to explore the space of all possible mixtures.

## 1. Introduction

### 1.1. Crop diversification with cultivar mixtures and overyielding

Besides the large increase in crop production due to the massive use of pesticides and fertilizers (Evenson and Gollin, 2003), the high-input farming system of industrialized countries comes at a cost. Indeed, pesticides were found to be harmful for the environment (IPBES, 2019), human health (Expertise collective, 2022), soil (Lal, 2007) and water (Moss, 2008), endangering food security in the long term. In contrast, crop diversification is promoted in the framework of the agroecological transition (Altieri, 1989; Gliessman, 2014) by means of different practices, such as cultivar mixtures for annual crops (Barot et al., 2017), with the goal of taking advantage of niche complementarity and facilitation (Brooker et al., 2015).

Cultivar mixtures, a first step towards crop diversification, are known to provide, on average, advantages in terms of disease reduction (Mundt, 2002; Wolfe, 1985). The benefits on yield are usually quantified with the comparison index named “relative yield total” (RYT) which equals the total yield of the mixture divided by the yield of the cultivars in pure stands weighted by their sowing proportions in the mixture (Reiss and Drinkwater, 2018; Weigelt and Jolliffe, 2003). For the case study of winter bread wheat, (Borg et al., 2018) showed in their meta-analysis of 32 studies that cultivar mixtures have a positive RYT on average, of +2% (i.e overyielding). However, they also showed that RYT displays a large variability (minimum RYT of -34% and maximum RYT of 50%, see figure 3). If the mechanisms driving disease control in mixtures are well studied (Borg et al., 2018; Pélissier et al., 2023), how cultivars interact in mixtures under non-limiting conditions for water, nitrogen, weeds and diseases is still understudied. Indeed, under such conditions, there still is a large variability in RYT between mixtures, for instance ±18% from the data of figure 3 in Forst *et al*. (2019). Our objective hence was to study the variability of RYT under conventional conditions, i.e. non-limiting for water, nitrogen, weeds and diseases, where light can still be a major limiting factor.

### 1.2. Differences in relative yields between mixtures and importance of plasticity

Deciphering the causes of RYT variability is therefore critical. First, wheat cultivars in pure stands are known to display a large variability for yield (grain weight per unit area), some outperforming the others in a given environment (Slafer et al., 2014). Moreover, the performance of any given cultivar can change when observed *between* stands, in pure vs. mixed stand, whether positively or negatively. This change in performance and related traits can be referred to as phenotypic plasticity, generally defined as the tendency for phenotypic variation to be influenced by the environment (Hallgrimsson et al., 2019). Regarding the large variability in RYT, the gaps between observed and expected mixture yields can hence be interpreted as phenotypic plasticity between pure and mixed stands with the change in neighboring plants between pure and mixed stands considered as the main environmental variable. Furthermore, a reaction norm being defined as “the systematic change in mean expression of a phenotypic character that occurs in response to a systematic change in an environmental variable” (De Jong, 1990), the difference in mean plant behavior between environments (pure versus mixed stands) for a given trait and cultivar hence constituted our reaction norm of interest. However, the extent of phenotypic plasticity in yield and its components, and the magnitude of plasticity differences between cultivars, remain largely unexplored, as well as the underlying traits and processes.

It is also imperative to go beyond the analysis of reaction norms for single traits separately given the importance of phenotypic integration that can lead to compensation between traits (Pigliucci and Preston, 2003). Beyond plant density (number of plants per m^2^), grain yield is decomposed into several components: spike density (number of spikes per m^2^), grain number per spike and thousand kernel weight, that display contrasted phenotypic plasticities (Sadras and Slafer, 2012). Among these components, a specific importance must be given to tillering as it is the most plastic trait in response to competition for light in winter bread wheat (Lecarpentier, 2017).

### 1.3. Objectives, hypotheses and design

To study the variability of RYT, our objectives in this paper were to assess the magnitude of phenotypic plasticity in pure versus mixed stands for yield and identify the yield components contributing the most to yield plasticity, the final aim being to link such plasticities to differences between cultivars in terms of functional traits, here height and earliness as these two traits were previously identified in Borg et al. (2018). For this we present an original field study of bread winter wheat, studying cultivar mixtures at plant scale, and thus enabling us to assess the above-ground plasticity of different life-history traits, whether as the ratio or difference of a given trait between in pure vs mixed stands. We focused on wheat because it is one of the most important crops in the world, representing 15% of cultivated areas, so any diversification effort will have a major impact.

First, we hypothesized (H1) that a cultivar’s plasticity in pure vs mixed stands was associated with the difference between its pure-stand yield and the mean of the pure-stand yields of the other cultivars with which it was mixed. This hypothesis stems from the “passive plasticity” definition (Forsman, 2015), where phenotypic changes are expected to be proportional to environmental differences. Here larger differences in cultivars’ traits in pure stands are expected to induce larger pure vs mixed stands changes (light, temperature, nutrients), leading to higher plasticity. Second, as the presence of neighbors is perceived early by plants, notably through light-quality signals (Demotes-Mainard et al., 2016; Smith, 2000), we hypothesized (H2) that differences in neighbor plant-plant interactions between pure and mixed stands were more likely to disproportionately affect the first yield component to be established, namely spike number, during the tillering dynamics. Third, as light quality is an unambiguous signal of canopy density (Smith, 2000), hence of above-ground biomass of neighbor plants, itself strongly correlated with plant height (Schirrmann et al., 2016), we hypothesized (H3) that height differences in mixtures during the crop cycle (rather than at maturity) were positively associated with the perception of light quality (early on) and quantity (later on), and hence correlated with plasticity in tillering dynamics.

A prerequisite to quantify such plasticities in yield and its components is the identification of each cultivar in the mixture, which is challenging and rarely done. We used cultivar-centered precision sowing to address this issue. In mixed stands, each plant of a given cultivar was sowed to be surrounded by plants of other cultivars, unlike mixtures in alternate ranks. To integrate a sufficient genetic diversity and make the continuous monitoring of individual plants feasible, notably their tillering dynamics in both pure and mixed stands, we designed an experiment with eight winter bread wheat cultivars. Moreover, we wanted our experiment to be representative of French farmers cultivating mixtures. The experiments were thus made under field conditions rather than in outdoor trays or inside a greenhouse. We also chose to study two quaternary mixtures as it is a typical mixture size used by French farmers (Mille et al., 2006). To contribute to the questions raised by (Borg et al., 2018) about the “need to quantify the effect of differences in height/earliness between varieties on overyielding and competitiveness”, two mixtures were assembled, one contrasted for earliness and the other for height.

## 2. Materials and Methods

### 2.1. Plant material

Among a panel of 210 European cultivars of winter bread wheat (Touzy et al., 2019), eight were chosen with contrasted heading dates and heights at maturity (all French cultivars except one German, Table S1): Accroc, Aubusson, Bagou, Belepi, Bergamo, Boregar, Expert and Kalahari. Two four-way mixtures were assembled, knowing that French farmers mix on average four to five cultivars (Mille et al., 2006). The first one, the earliness mixture, was composed of Accroc, Aubusson, Bergamo and Expert, the first two having earlier heading dates than the two others. The second one, the height mixture, was composed of Bagou, Belepi, Boregar and Kalahari, with Belepi and Kalahari known to be taller at maturity. Both mixtures were assembled in equi-proportions, i.e the proportion of each cultivar at sowing was 25%.

### 2.2. Field trial

A field trial was conducted at Le Moulon, Gif-sur-Yvette, France (latitude 48.710192, longitude 2.160042, altitude 161m) on deep loamy soil, from sowing on October 30, 2019 to harvest on July 20, 2020. It was fully replicated once, with sowing on November 5, 2020, and harvest on August 2, 2021. Each year harvest was performed after the latest cultivar achieved maturity. The differences in earliness and at maturity were lower than those at heading, and were not an issue at harvest. Each year, the whole trial was a rectangle of 8m wide and 12m long, and comprised 12 “nano-plots” (0.88m x 2.32m = 2m^2^ for pure stands and 0.88m x 1.68m = 1.5m^2^ for mixed stands): 8 pure stands and 4 mixed stands (two replicates per mixture). All nano-plots were sown as a regular lattice made of 12 ranks, with 30 rows for pure stands and 22 rows for mixed stands (Figure S1). The pure stands had more rows to allow for destructive sampling all along the crop cycle. Both row and rank spacings measured 8 cm corresponding to a sowing density of 160 seeds.m^-2^, allowing both a strong interaction between plants, and sufficient tillering plasticity (multiple tillers per plant). The areas of the nano-plots, similarly to other studies (Chen et al., 2021), were chosen to allow for the non-destructive phenotyping of individual plants in the center of the plots without damaging other plants. The spatial distribution of each mixture (Figure S1) was obtained by randomizing each cultivar (Lieng et al., 2012), and seeds were sown at the center of PVC rings of 0.2cm thickness, 5.5cm radius and 5cm depth (Figure S2) to facilitate the phenotyping and harvest of individual plants without disturbing their growth. In terms of agronomic management, seeds were treated prior to sowing with Celest Extra, and fungicides were applied during the crop cycle: the first year 250 L.ha^-1^ of Sportak + 1 L.ha^-1^ of Tebucur on 2020/04/03, 2 L.ha^-1^ of Rivexo on 2020/05/18 and 1 L.ha^-1^ of Caramba Star on 2020/05/29, and the second year 2 L.ha^-1^ of Rivexo on 2021/05/12 and 1 L.ha^-1^ of Caramba Star on 2021/06/10. Nitrogen fertilization was applied twice, at the BBCH30 and BBCH55 stages, with respectively 70 and 40 kg.ha^-1^. The preceding crop for the 2019- 2020 trial was a fallow, and it was alfalfa for the 2020-2021 trial. Corridors between nano-plots were regularly weeded by hand. Average daily temperature, rainfall and global radiation were measured by a local weather station and monitoring data was retrieved from the INRAE CLIMATIK platform (https://agroclim.inrae.fr/climatik/, in French) managed by the AgroClim laboratory of Avignon, France. The average daily temperature between October and August was 12.1°C in 2019-2020 and 10.8°C in 2020-2021, cumulative rainfall was 617 mm in 2019- 2020 and 665 mm in 2020-2021, and cumulative global radiation was 4261.22 MJ.m^-2^ in 2019-2020 and 3447.19 MJ.m^-2^ in 2020-2021.

### 2.3. Phenotyping

For each year, from January until June, multiple above-ground traits were phenotyped once a month for a total of six time points. In pure stands, 36 plants were destructively sampled at each time point to record height (PH, in cm), tiller number (TN) and above-ground dry biomass (PW, in g) after a 48h-drying at 60°C. In mixed stands, height and tiller number were recorded non-destructively at each time point for each individual plant. Phenological stages BBCH30 (ear at 1cm) and BBCH55 (half of ear emerged above the flag leaf ligule; heading) were determined based on plants in pure stands (Table S2). At harvest all remaining plants were sampled to record the following variables at the individual-plant scale: for plant *i*, shoot dry weight (SW_i_, in g), spike dry weight (SPW_i_, in g), spike number (SN_i_), grain number (GN_i_) and grain weight (GW_i_). From these were then computed the grain number per spike (GNpS_i_ = GN_i_/ SN_i_) and the thousand kernel weight (TKW_i_ = GW_i_ / GN_i_ * 1000, in g). To account for border effects, the data from all plants sown along a border of a nano-plot, whether in pure or mixed stand, were discarded, i.e only the data from inner plants were analyzed. As a result, at a given time point, the data from 20 (respectively 59 to 90) plants were analyzed for each cultivar in pure (resp. mixed) stand. In June 2020 most plants in the Aubusson pure stand were destroyed by coypus before harvest. Missing data for SW, SPW, GN and GW were imputed from another field experiment performed the same year at a neighboring site, using the multivariate model implemented in R package mice (Buuren and Groothuis-Oudshoorn, 2011).

### 2.4. Statistical analysis

#### 2.4.1. Replicates of mixtures

Each year, each mixture was cultivated in two replicate nano-plots. For each year separately, the effect of the replicate, α_r_, was assessed per cultivar in each mixture with an ANOVA for each of the phenotyped variables (PH and TN per time points; SW, SPW, SN, GN, GW, PW, HI, GNpS and TKW at harvest): for each plant *i* in replicate *r*, its phenotypic value y_ir_ was modeled as y_ir_ = μ + α_r_ + ε_ir_ with ε_ir_ ∼ N(0, σ^2^). As such replicate effects were not significant at 5% (type I error rate) in the vast majority of the cases (data not shown), the following analyses were done using the data from both replicates without a replicate effect.

#### 2.4.2. Yield

The yield of a given cultivar, whether in pure or mixed stand, was computed by dividing the mean grain weight per plant of this cultivar by the area of one plant in the stand computed as the inverse of the sowing density (1/160=0.00625 m^2^). For French official variety trials, called Value for Cultivation, Use and Sustainability (VCUS) testing, the GEVES accounts for border effects using inter-row and inter-plot distances which typically increases the microplot area used for yield calculation by 20%. Hence in our case, in addition to discarding the data from border plants, the area of one (inner) plant corrected for borders equaled 0.00750 m^2^.

#### 2.4.3. Relative yield totals

The relative yield total of a given mixed stand (Reiss and Drinkwater, 2018) was computed as RYT = Y_mix_ / (p_1_ × Y_pure,1_ + p_2_ × Y_pure,2_ + p_3_ × Y_pure,3_ + p_4_ × Y_pure,4_) where Y_mix_ was the grain weight per plant averaged over all plants in the mixed stand, p_c_ was the proportion of plants in the mixed stand being of cultivar *c* in the mixed stand, and Y_pure,c_ was the grain weight per plant averaged over all plants in the pure stand of the given cultivar. A value above 1 (respectively below 1) indicates overyielding (respectively underyielding). The RYT was also expressed in percentage: 100 ×(RYT-1), so that a positive value (respectively negative) indicates overyielding (resp. underyielding).

#### 2.4.4. Cultivar rankings according to dominance

A cultivar *c* will be said to *dominate* a cultivar *c’* within a given stand when the plant grain weight of *c* is higher than the one of *c’*, either if observed in parallel pure plots, or in the same mixture plot. Dominance rankings between cultivars of a given mixture were defined based on Tukey’s range test on average plant grain weight using the R package multcomp (Hothorn et al., 2008). This test was applied per mixture and year. Pairwise differences between cultivars were declared significant if their p-values were below 0.05, followed by a letter-based representation (Piepho, 2004). Dominance rankings were established in ascending order based on this letter representation and decimal ranks were given to cultivars belonging to two groups. The same procedure was also applied on the pure-stand data for all cultivars of a given mixture.

#### 2.4.5. Tests of phenotypic plasticity (reaction norms)

For each cultivar per year, the equality of means of different traits between pure and mixed stands was tested. Our first index of phenotypic plasticity, PPI_1_(X), hence was the difference between the mean value of a trait X in mixed (μ_mixed_) vs pure (μ_pure_) stands: PPI_1_(X) = μ_mixed_ – μ_pure_. The null hypothesis was that the mean of the trait in the pure stand was equal to its mean in the mixed stand, meaning no plasticity in mixtures. Graphically, a line can be drawn from μ_pure_ to μ_mixed_ (Figure 1) and corresponds to what is called a (linear) “reaction norm” (Guntrip and Sibly, 1998). Moreover, the sign and significance of the slope of a reaction norm are the same as the sign and significance of PPI_1_. The hypothesis was tested separately for different traits. A bootstrap algorithm was applied, assuming neither equal variances between pure and mixed stands nor the normality of trait values per stand. One thousand bootstrap samples were generated to approximate the distribution of the test statistic under the null hypothesis, and a p-value was computed based on this distribution. No correction was applied to correct for multiple testing over all traits and years as justified in an exploratory context (Heller, 2011; Rothman, 1990).

**Figure 1:**
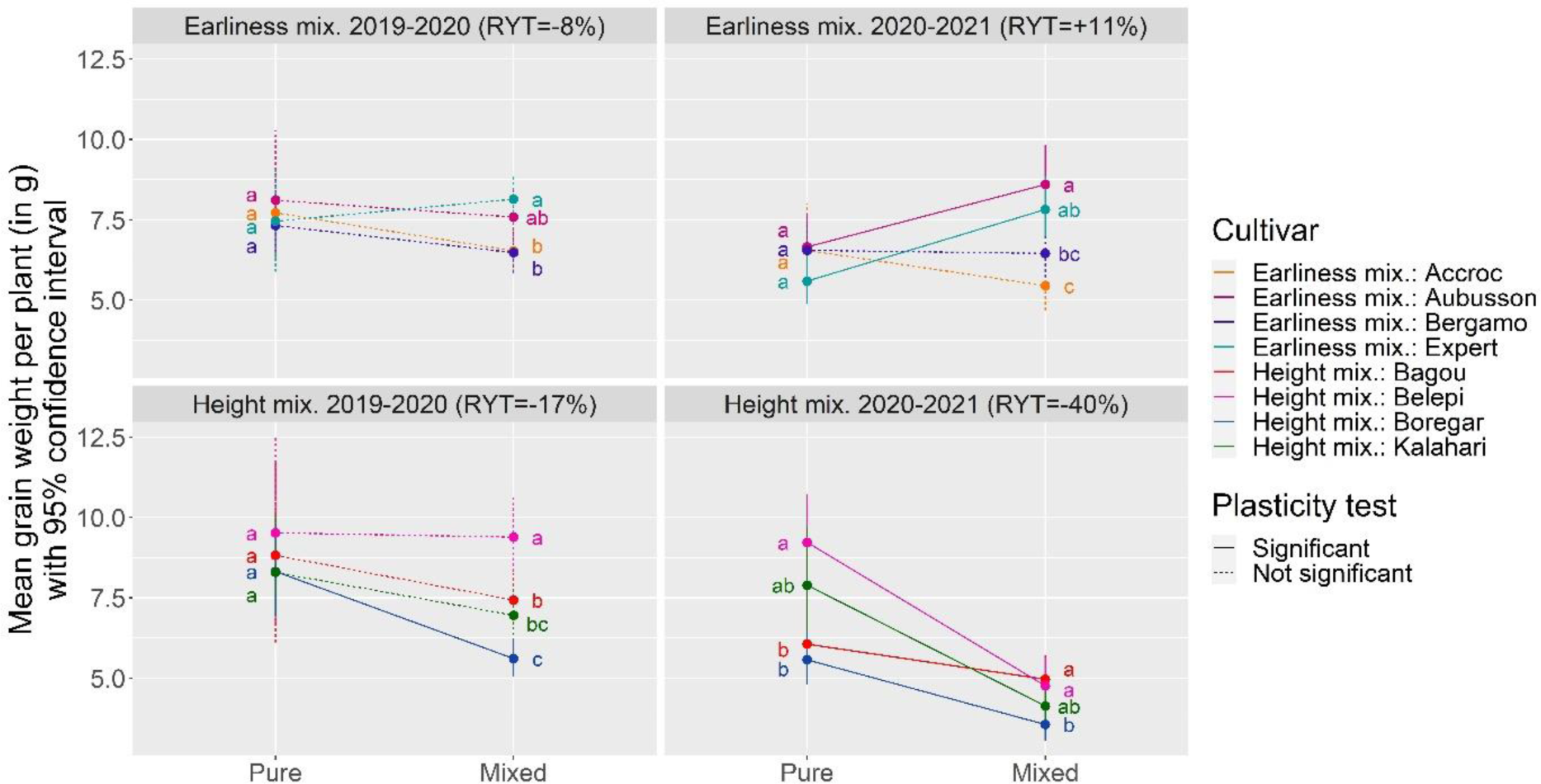
Reaction norms of mean grain weight per plant between pure and mixed stands per cultivar for both mixtures in both years. Vertical segments represent confidence intervals at 95%. Letters indicate pairwise differences between cultivars in pure or mixed stands. Line types indicate if the plasticity index PPI_1_ (difference between the mean value of grain weight per plant in mixed vs pure stands) is significantly different from zero or not.

#### 2.4.6. Test of the relation between plant grain weight plasticity and plant grain weight difference between neighbors

As a second index of phenotypic plasticity PPI_2_, the plant grain weight plasticity of a given cultivar *f* (considered as focal) in a given year was quantified using the opposite of the relative competition intensity (fifth index indicated as nr. 3 in table 1 of (Weigelt and Jolliffe, 2003), i.e., PPI_2_(GW_f_) = 100 x (GW_mix,f_ - GW_pure,f_) / GW_pure,f_, where GW_s,f_ is the mean grain weight of the focal cultivar *f* in stand *s* (pure or mixed) in the given year. The same year, the relative plant grain weight difference between the focal cultivar and its neighbor cultivars, noted *ΔGW*_*pure,f,y*_, was quantified as the pure-stand plant grain weight of the focal cultivar *f* in comparison with the mean of the pure-stand plant grain weights of its three neighbor cultivars in the mixture, noted mGWpure,f, with {*n* ∊ G \ *f*} being the set of neighbors cultivars and G the set of all cultivars in the mixture:

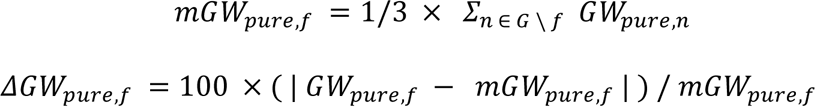

The absolute value of the second plasticity index PPI2(GW) was then regressed on the absolute value of ΔGWpure,f, per year, using a linear model:

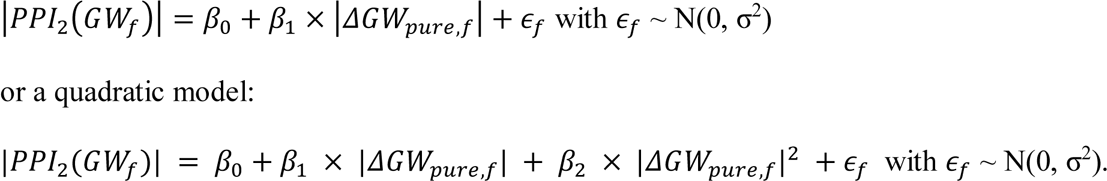

Both models were compared per year using the likelihood ratio test as computed by the anova function in R.

#### 2.4.7. Decomposition of plant grain weight plasticity into the plasticity of its components

The grain weight of a plant *i*, GW_i_, can be decomposed into the product of its spike number (SN_i_), its grain number per spike (GNpS_i_) and its single kernel weight (SKW_i_) as SN_i_ × GNpS_i_ × SKW_i_ with SKW_i_ = TKW_i_/1000. For a given cultivar in a given year, we started by summing each variable over all n_s_ harvested plants (s indicating the stand, pure or mixed) to get X_s,tot_ where X is one of GW, SN and GN, and we then computed GNpS_s,tot_ = GN_s,tot_ / SN_s,tot_ and SKW_s,tot_ = GW_s,tot_ / GN_s,tot_, where SN_s,tot_ is the average of spike number over al n_s_ harvested plants. As a third index of phenotypic plasticity (PPI_3_) between pure and mixed stands for a given trait of a given cultivar in a given year, we used the natural log of the ratio of the trait value in mixture X_m,tot_ over the trait value in pure stand X_p,tot_ such as PPI_3_(X)=log(X_m,tot_/X_p,tot_). This index was computed for GW, SN, GNpS and SKW. Thanks to the property of the log turning multiplication into addition, the index for plant grain weight PPI_3_(GW) can be decomposed as the sum PPI_3_(SN) + PPI_3_(GNpS) + PPI_3_(SKW). The contribution of each component plasticity to plant grain weight plasticity were compared based on their absolute value. As an example, |PPI_3_(X_1_)| > |PPI_3_(X_2_)| was interpreted as trait X_1_ plasticity contributing more than trait X_2_ plasticity to grain weight plasticity. Moreover, note the non-linearity of the log, so that an absence of plant grain weight plasticity, PPI_3_(GW)=0 may be due to PPI_3_(SN)=- 1, PPI_3_(GNpS)=+1 and PPI_3_(SKW)=0, in which the reduction of 63% of SN in mixture compared to pure stand is compensated by an increase of 172% of GNpS.

#### 2.4.8. Plasticity in tiller emission and regression

The numbe of spikes of a given plant (SN) was decomposed into a sum of two metrics, SN = MTN + (-NRT) where MTN was the maximal number of tillers and NRT the number of regressed tillers. The equality of means of each metric between pure and mixed stands was tested for each cultivar and year separately with the same bootstrap approach as for testing phenotypic plasticity with PPI_1_ above (see 2.4.5.).

#### 2.4.9. Correlation between tiller plasticity and height differential in mixture

For each year *y*, the height differential in mixture between cultivar *c* and the other cultivars was computed at each date *d* (in days after sowing) during tiller cessation (between sowing and the end of tiller emission): ΔPH_yc,mix_(d) = PH_yc,mix_(d) - (∑ _j ≠ c_ PH_yj,mix_(d)) / 3. The height differential during tiller emission, when the maximal number of tillers (MTN) is reached, was compared with the second plasticity index in pure versus mixed stands for the maximal number of tillers (MTN) computed as PPI_2_(MTN). Similarly, the height differential was also computed at each date *d* during tiller regression (between tiller cessation and maturity). This index of height differential during tiller regression was compared with the second plasticity index in pure versus mixed stands for the number of regressed tillers (NRT) computed as PPI_2_(NRT). For the analysis of the tiller-regression phase, we fitted three linear models, all with the same response variable PPI_2_(NRT), but differing in their explanatory variables: the first model with only the height differential, the second model adding the plasticity index for tiller emission PPI_2_(MTN), and the third model with also the interaction between both explanatory factors. The best model was selected according to a likelihood ratio test.

#### 2.4.10. Reproducibility

Analyses were conducted in R (R Core Team, 2020) and figures were produced with the ggplot2 package (Wickham, 2009). Supplementary information, data and code that support the findings of this study are openly available at the INRAE space of the *Recherche Data Gouv* repository https://doi.org/10.57745/LZS8SU.

## 3. Results

### 3.1. Contrasted yields between the two years

The average yield in our field trials of winter bread wheat (see subsection 2.4.2) equaled 9.3 t.ha^-1^ (11.15 t.ha^-1^ before correction for border effects), and it was associated with an average ear density of 593 spikes.m^-2^. The first year (10.31 t.ha^-1^; 630 spikes.m^-2^) was more productive than the second one (8.32 t.ha^-1^; 555 spikes.m^-2^). Looking at pure stands, this yield decrease was especially strong for all four cultivars of the earliness mixture (-17%) and two cultivars (Bagou, Boregar) of the height mixture (-32%). Regarding mixed stands, the total yield of the height mixture sharply decreased the second year compared to the first (-41%) whereas the yield of earliness mixture remained stable (-1%). The lower yields in the second year were associated with lower global radiations starting after BBCH30 as well as lower rainfall between BBCH30 and BBCH77 and higher rainfall after BBCH77 (Figure S3).

### 3.2. Significant plant grain weight plasticity in pure versus mixed stands led to dominance relationships in mixtures

In the four mixture-year combinations, the plant grain weight observed in the mixture deviated from the expectation based on the plant grain weights in pure stands and the sowing proportions, as shown by the reaction norms between pure and mixed stands (Figure 1). The earliness mixture performed globally well, with a positive Relative Yield Total (RYT) the second year (+11%, -8% the first year, with an ear density of 577 spikes.m^-2^ both years), whereas the height mixture was underyielding both years (RYT=-17% and -40%, with 687 spikes.m^-2^ and 529 spikes.m^-2^, in 2019-2020 and 2020-2021 respectively).

Dominance relationships among cultivars were defined by testing pairwise plant grain weight differences and turning them into rankings (see subsection 2.4.4 and details in Table S3), i.e. a genotype is said to be dominating another if its plant grain weight is higher. Although trivially done for the pure stands, this was made possible in the mixed stands thanks to our experimental design in which the cultivar of each plant is known. For three mixture-year combinations out of four, there was no dominance among cultivars in pure stands (Figure 1). In contrast in mixed stands, all mixture-year combinations revealed ranking hierarchies between cultivars, i.e. dominance relationships. Noticeably, the two most dominant cultivars were stable over both years (Aubusson and Expert in the earliness mixture, Bagou and Belepi in the height mixture). They were not necessarily the tallest nor the earliest cultivars in their respective mixtures.

Accordingly, neither dominance rankings nor mean plant grain weights were significantly correlated between pure and mixed stands (Figure S4).

Measuring plant grain weight in mixtures per cultivar also allowed us to compute a first index of phenotypic plasticity (PPI_1_) between pure and mixed stands for each cultivar (see subsection 2.4.5) and test its significance. In Figure 1, PPI_1_ was represented as a line (linear reaction norm) whose type indicate the significance of the test. Out of the sixteen cultivar-year combinations, 7 (44%) displayed a significant plasticity. Among these, two had positive reaction norms, Aubusson and Expert in the earliness mixture in 2020-2021, hence explaining the overyielding of this mixture.

The magnitude of plant grain weight plasticity in pure versus mixed stand for a given cultivar, using a second phenotypic plasticity index |PPI_2_| was regressed on the absolute difference between its grain weight in pure stand and the average grain weight of the other cultivars in pure stand with which it was mixed (see subsection 2.4.6 and Figure 2). The association in the first year was not significant at α=5% (slope p-value = 0.35). The second year however, the association was significant at α=5% (slope p-value = 0.046 and R^2^ = 0.51) with a quantile-quantile plot of residuals being reasonable given the small sample size. The second year, even though a quadratic model could provide a fit visually more appealing, the linear model was retained based on the likelihood ratio test (p-value = 0.21). Note that after removing the right- most point (Belepi in 2020-2021), this relation is just above the significance threshold (p-value = 0.0563).

**Figure 2:**
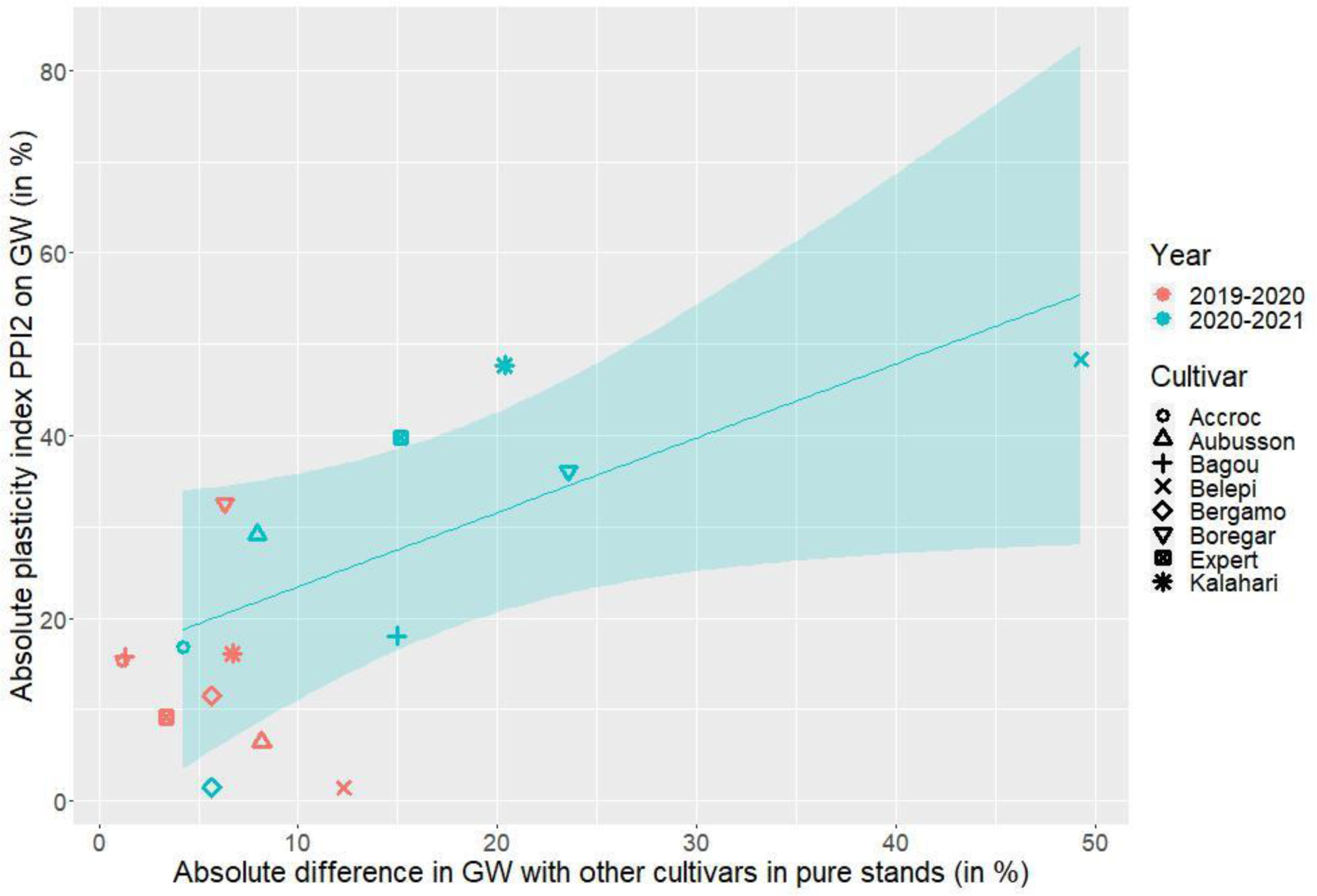
Plasticity index |PPI_2_| (difference between the mean value of grain weight per plant in mixed vs pure stands, relative to pure-stand plant grain weight, in percentage) between pure and mixed stands for plant grain weight (GW) per cultivar and year versus the absolute difference between the pure-stand plant grain weight of the focal cultivar and the mean of the pure-stand grain weights of the other cultivars in the mixture. Only significant regression lines are represented.

**Figure 3:**
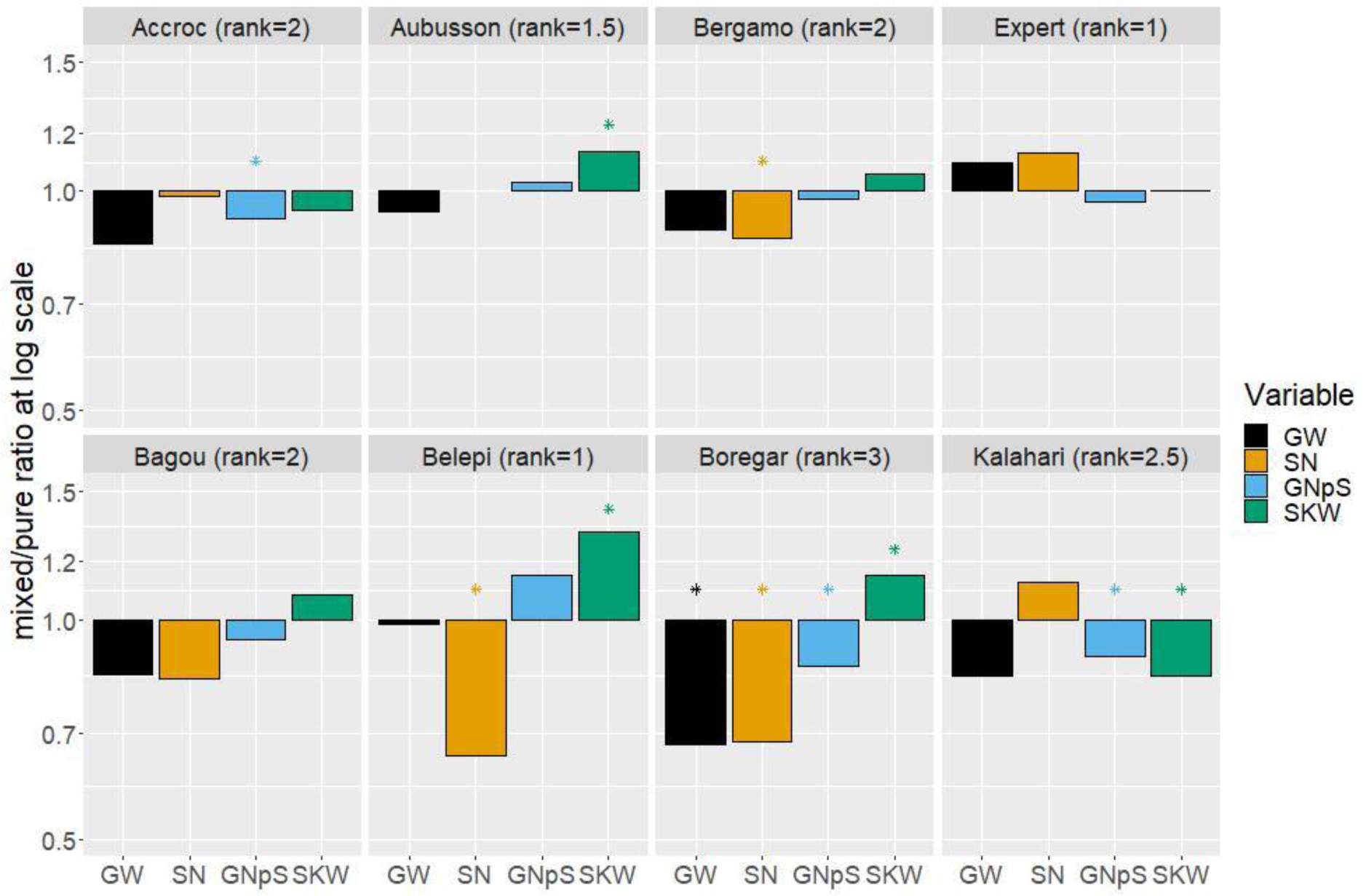
Values of phenotypic plasticity (as the ratio mixed/pure) in log scale of plant grain weight (GW) and its components, spike number (SN), grain number per spike (GNpS) and single kernel weight (SKW) in 2019-2020. Note that the y-axis is in log scale, allowing for the additivity of the components. Stars indicate significant differences in pure versus mixed stands for a given variable, cultivar and year. Note the missing data for Aubusson due to coypus predation in pure stand.

### 3.3. Plasticity in spike number explained most of grain weight plasticity

Given the significant phenotypic plasticity in grain weight displayed by some cultivars, we proposed a new way to decompose this plasticity into the phenotypic plasticity of its components using a third index PPI_3_ (see subsection 2.4.7, Figure 3 for 2019-2020, Figure S5 for 2020-2021). Out of the sixteen cultivar-year combinations, the seven with significant grain weight plasticity always had a significant plasticity for at least one of its components. Moreover, among the nine other combinations with no grain weight plasticity, five had at least one plastic component. Overall, the plasticity in spike numbers was the main contributor, in five out of seven cases in 2019-2020 (notably for three cultivars in the height mixture), and also six out of eight cases in 2020-2021. Moreover, the plasticities in grain weight components were strongly correlated, although with a change of sign between years (Figure S6).

### 3.4. Plasticity in both tiller emission and regression explained plasticity in spike number

Spike number is the result of tillering, a dynamic process whose particularity is to be non-monotonic. Thanks to our experimental design, we were able to phenotype this process for the first time at the level of individual plants in field conditions for both pure and mixed stands, as exemplified for the height mixture in 2019-2020 (Figure 4; see Figure S7 for the other cases).

**Figure 4:**
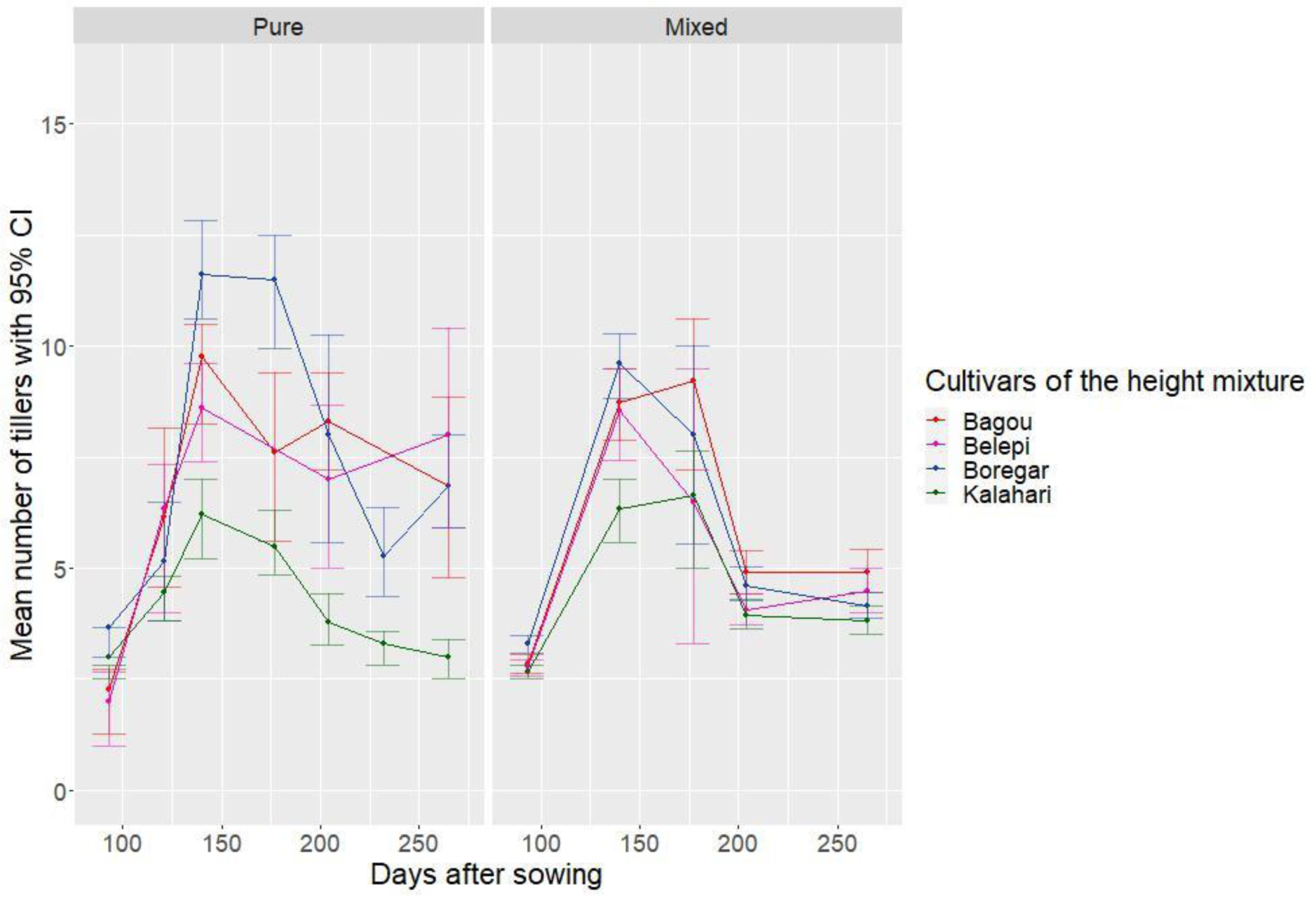
Average tillering dynamics from individual plant data all along the crop cycle for each cultivar of the height mixture in 2019-2020 for both pure and mixed stands. Error bars correspond to 95% confidence intervals.

In qualitative terms, the two phases of tillering dynamics, emission and regression separated by a plateau, were clearly visible for both pure and mixed stands. However, these dynamics looked quantitatively different between both stands. Beyond their outcome, spike number, the tillering dynamics themselves were also plastic. Indeed, our two metrics, the maximum number of tillers (MTN) for the emission phase and the number of regressed tillers (NRT) for the regression phase, showed several significant cases of plasticity between stands, as exemplified for the earliness mixture in 2019-2020 (see subsection 2.4.8, Figure 5 for the height mixture in 2019-2020 and Figure S8 for the other cases). Furthermore, the plasticities of these metrics could compensate for each other (Figure 5), sometimes completely, hence leading to no plasticity in the spike number, or only partially hence leading to spike number plasticity. Reaction norms for the proportion of regressed tillers (NRT/MTN) is available in supplementary material (Figure S9).

**Figure 5:**
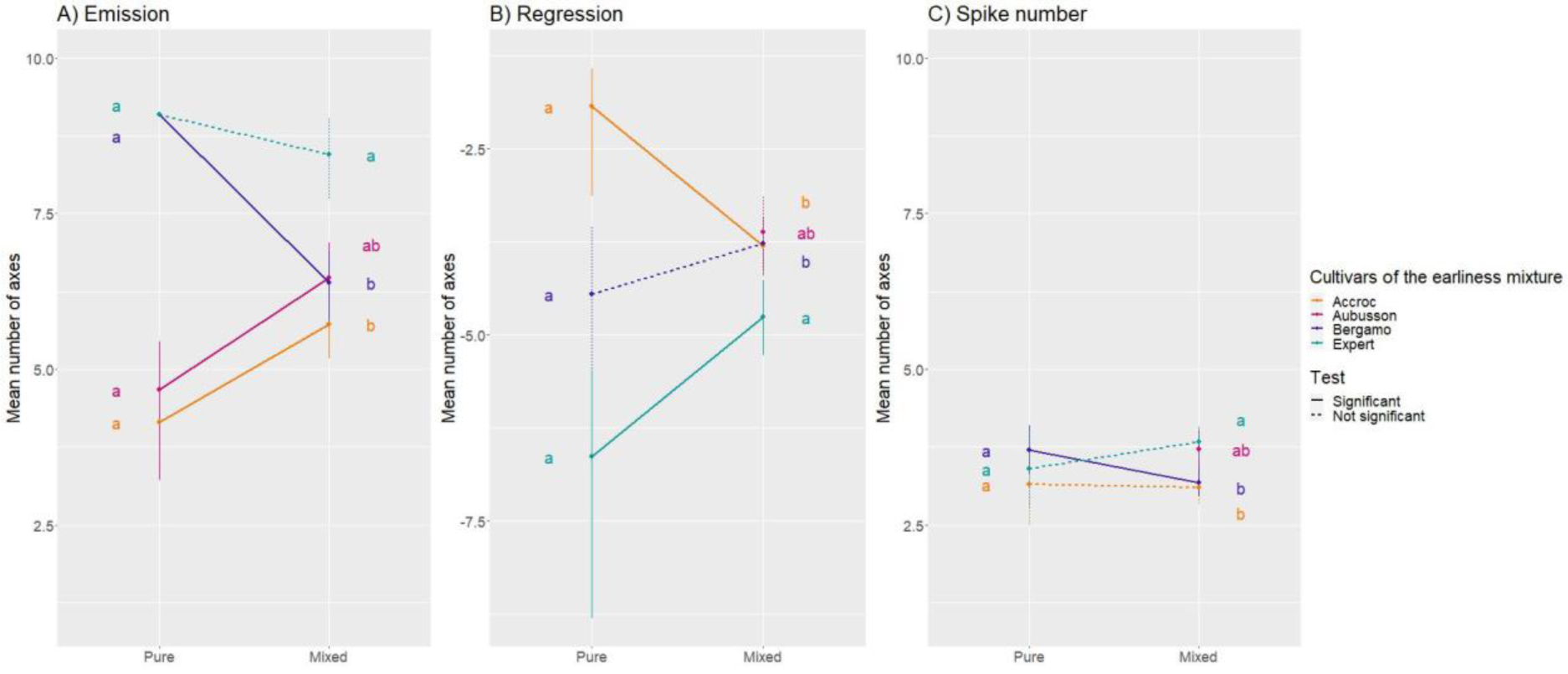
Reaction norms of number of axes per plant during tiller emission (MTN) and regression (-NRT) between pure and mixed stands per cultivar for the earliness mixture in 2019-2020. Vertical segments represent confidence intervals at 95%. Letters indicate pairwise differences between cultivars in pure or mixed stands. Line types indicate the statistical significance of the reaction norms. Note the two missing missing data in subplots B and C for Aubusson in pure stand.

### 3.5. Dynamics of height differentials explain tillering plasticity

Knowing that tillering dynamics are influenced by the perception of a light-quality signal, we monitored the height dynamics of individual plants (Figure 6 and Figure S10), testing for phenotypic plasticity between pure and mixed stands at each time point (Figure S11).

**Figure 6:**
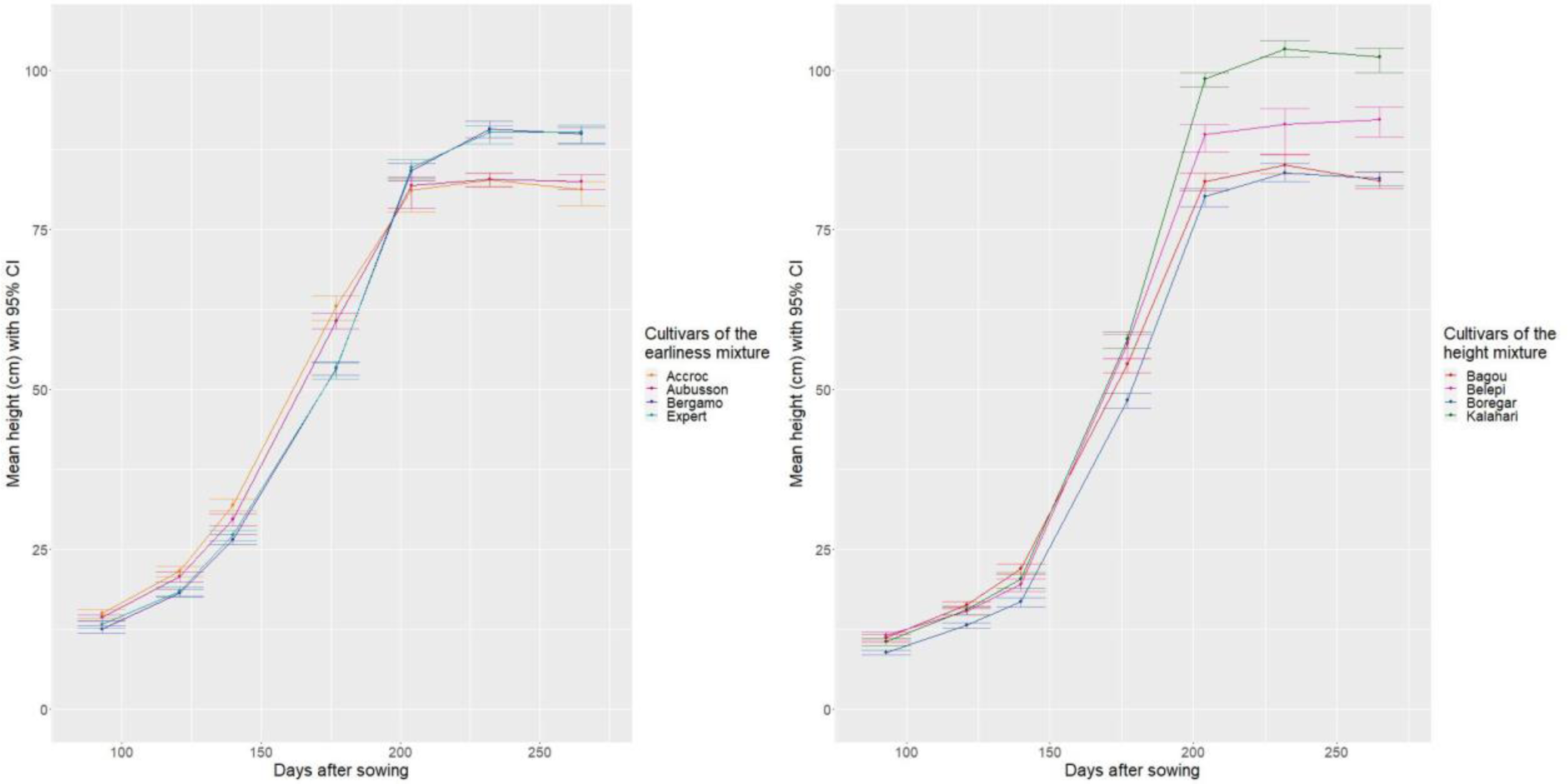
Mean plant height (in cm) throughout growth for the different cultivars grown in mixed stands for both mixtures in 2019-2020. Vertical segments represent confidence intervals at 95%.

In the earliness mixture in 2019-2020, the two cultivars chosen for their earliness, Accroc and Aubusson, were indeed taller than their neighbors, Bergamo and Expert, early in the crop cycle because the elongation of their main stem started earlier. In the height mixture in 2019-2020 the two cultivars chosen for their higher height at maturity, Belepi and Kalahari, were indeed taller at the end of the growth cycle.

Moreover, during the phase of tiller emission, the height differentials per cultivar (in comparison with the others) in the mixtures were significantly associated with their plasticity in maximum tiller number (see subsection 2.4.9 and Figure 7). The slope of the linear regression was positive for both years. We also tested this relation during the phase of tiller regression (Figure S12) while also controlling for the plasticity during the phase of tiller emission. The plasticity in tiller regression was significantly associated with both height differential and plasticity on tiller emission only in 2020-2021.

**Figure 7:**
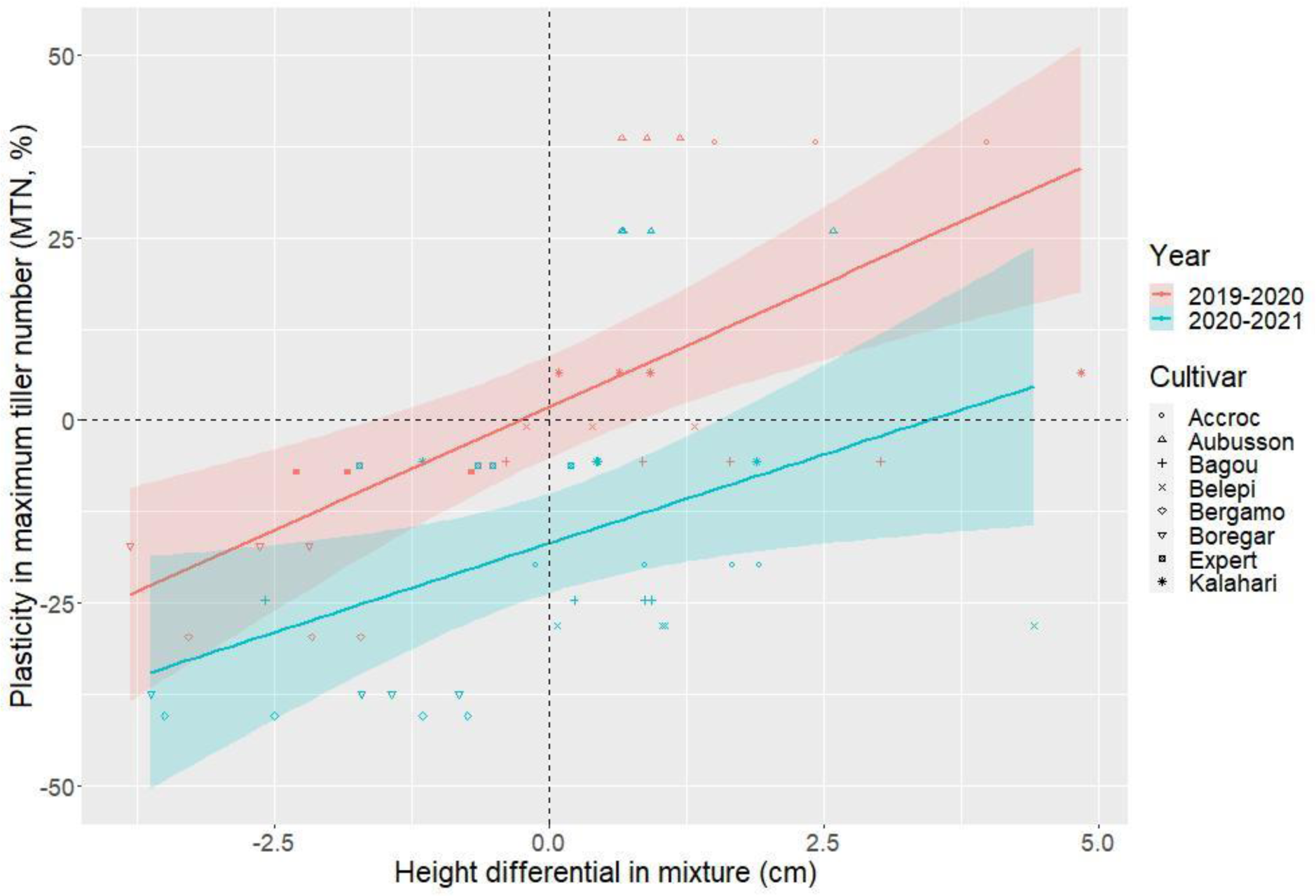
Plasticity of the maximum tiller number (MTN) versus height differentials in mixture computed during the tiller emission phase. Both significant regression lines are displayed, here slope = 6.742 ± 1.581 in 2019-2020 (pvalue 2.69×10^-4^, R^2^=0.43) and slope = 4.870 ± 1.957 in 2020-2021 (p-value 0.0186, R^2^=0.17).

## 4. Discussion

We proposed a new experimental design and analytical framework to study, in field trials, the reaction norms of cereals in pure versus mixed stands. We applied them to scrutinize the development and productivity of eight bread wheat cultivars, in pure stand and into two quaternary cultivar mixtures, and replicated the trial in two different years. The main advantage of the new design consisted in giving access to the behavior of each individual plant, and hence each cultivar in a mixture. As a result, it allowed the study of yield phenotypic plasticity per cultivar between pure and mixed stands, as exemplified by our results using three indices of phenotypic plasticity. We notably proposed a new index to study such a plasticity by decomposing it into the plasticities of its components. Although our design and framework came with limits discussed below, they delivered new results summarized right after.

### 4.1. Limits of the experimental design

Due to the need to know the cultivar of each individual plant in the mixtures and to continuously monitor them over time, we used PVC rings in mixtures to better separate the plants and facilitate measurements, whereas we performed destructive measurements in pure stands. This could have introduced a bias in mixture measurements, which we considered negligible although we did not assess it. Such a design can also be of interest to study intercropping, as long as the plants are easily separable to be phenotyped individually. This should be the case of mixtures of cereals (McAlvay et al., 2022), but not of some cereal-legume mixtures such as those with vining cultivars of field pea whose tendrils coil around neighboring plants, allowing them to climb and use them as support (Podgórska-Lesiak and Sobkowicz, 2013).

As motivated in the introduction, by applying nitrogen fertilization as well as fungicides, and by removing weeds manually, we focused our experiments on conditions under which light availability was the main limiting resource underlying competition between wheat plants. Water stress was also not deemed impactful enough to require irrigation. Yet, abiotic and biotic factors have an impact on the outcome of plant-plant interactions. Resources such as nutrients and water are known to trigger competition mechanisms other than those for light (Craine and Dybzinski, 2013). As a consequence, the RYTs of specific cultivar mixtures may change from over to underyielding, and inversely, depending on the timing and intensity of such stress factors. Moreover some crop traits may be advantageous to limit weed growth (Lazzaro et al., 2019; Wicks et al., 2004) but, as a corollary, they may also limit the growth of other cultivars in a mixture. Overall, our results may change under strong abiotic or biotic stress, but the proposed framework could nevertheless quantify plasticity in yield components under any stress scenario.

Among the studied yield components, spike number results from tillering dynamics, known to be strongly dependent on plant density (Darwinkel, 1978). We hence chose for our field trials a density of 160 plants.m^-2^ allowing strong interactions between plants during the tillering process while maintaining a sufficient number of spikes per plant at harvest (3.57 on average over the two years). This density is slightly lower than the range advised to farmers in central and northern France (170-400 plants.m^-2^; Arvalis pers. com). Nevertheless, other plant densities are likely to have an impact on the magnitude and sign of phenotypic plasticities and this impact is worth investigating further, as well as the effect of inter-row distance (Abichou et al., 2019). Finally, we chose eight commercial cultivars (“elite”) to compose two quaternary mixtures but the use of more diverse cultivars (eg. landraces) and other mixtures of other orders (eg. binary) may produce different results. Besides, the choice of cultivars was based on earliness and height at maturity, but cultivars of the earliness mixture also showed differences in height at maturity (and vice-versa cultivars of the height mixture also showed differences in earliness) as there were no perfect combinations in the panel of available cultivars, but these differences were not as important.

Our choices regarding cultivars, mixture size and management were made so that plant biomass production and grain allocation would be representative of what farmers in northern and central France do and experience. Notably, yields from the field experiment were in the range of yields from farmers’ fields in northern and central France (Brisson et al., 2010).

### 4.2. Significant reaction norms of yield components between pure and mixed stands

Plant grain weight displayed a large variation between pure and mixed stands. Over the four mixture-year combinations, only one favorable case (RYT>1) was observed, contradicting the usual overyielding of mixtures on average (Borg et al., 2018). The observed variation in RYT was due to the phenotypic plasticity of cultivars, with both positive and negative reaction norms from pure to mixed stand that affected the dominance relationships between cultivars. The results also confirmed our first hypothesis: the second year, the magnitude of plant grain weight plasticity for a given cultivar was significantly positively associated with the difference of its plant grain weight with the three other cultivars, observed in pure stand. The fact that this association was not significant the first year could be explained by the better yield potential of the first year weather conditions, leading to much smaller yield differences between cultivars in pure stands, in line with the stress gradient hypothesis (Malkinson and Tielbörger, 2010). Nevertheless, the yield difference among cultivars in pure stands can be regarded as an indicator of asynchronous production trends, also called asynchrony, a notion with increasing interest for the assembly of cultivars in mixtures (Stefan et al., 2024; Valencia et al., 2020; Weih et al., 2021). Thanks to a new decomposition, we then showed that plant grain weight plasticity resulted from the plasticities of its component that can cancel or amplify each other. We confirmed our second hypothesis of a main contribution of spike number plasticity, similarly to what was observed in rice (Jennings and Aquino, 1968). Plasticity in spike number was itself driven by plasticity in both the tiller emission and regression phases that, importantly, could compensate for each other. In agreement with our third hypothesis, the height differentials of cultivars in mixture were significantly associated with the tillering plasticities, notably for the maximal number of tillers. Finally, our results highlighted the importance of the environmental conditions, with several relationships changing sign from one year to the next.

Overall our results showed that the most integrative trait of interest, plant grain weight, can be stabilized in mixture over contrasted years, as well as its main component, spike number, through the net result of cancelation-amplification of the underlying determinants.

### 4.3. Plasticity in tillering

In general, phenotypic plasticity of a given trait can be classified as advantageous, neutral or disadvantageous depending on the relationships with fitness (Alpert and Simms, 2002). Moreover, the plasticity in a given trait can be qualified as *active* in the sense of “anticipatory, and often highly integrated, phenotypic changes in response to some environmental cue or signal”, whereas the plasticity in another trait would be best viewed as *passive*, i.e., as a “consequence of the environment, such as stunted growth owing to low resource levels” (Forsman, 2015). In the case of small-grain cereals in general, and wheat in particular, the phenotypic plasticities of yield and its components have been widely studied, mostly in pure stands (Sadras and Rebetzke, 2013; Sadras and Slafer, 2012; Slafer et al., 2014), but without discussing their active versus passive nature. As a consequence, little is known in mixed stands about either the impact of competition between cultivars on these plasticities, or the other way around, i.e the impact of these plasticities on the outcomes of the competition between cultivars. Moreover, a discussion of active versus passive plasticity of each yield component is lacking, especially in heterogeneous stands such as cultivar mixtures for which our study specifically contributes.

Given the early detection of light signals (R:FR), their integration as a bet on future competition (Schmitt and Wulff, 1993) and the consequences in terms of architectural changes (Casal, 1988; Evers et al., 2006; Xie et al., 2016), plasticity in tillering can be characterized as active. As the outcome for a plant of an early bet, here on tillering, can be more consequential on plant biomass, grain weight (and overall fitness) than would be a late bet, such as on thousand kernel weight, improving the odds of the bet on tillering would explain that evolution selected for the capacity to detect variation in light cues and respond appropriately to them. Moreover, we observed a hierarchy of yield-component plasticities in pure versus mixed stands in agreement with known results in the literature obtained when comparing, in pure stand, stress versus no- stress conditions (Sadras and Slafer, 2012). In the majority of cases, the contribution of the plasticity in spike number to plant grain weight plasticity was higher than the ones from the plasticities of grain number per spike and thousand kernel weight. The plasticity hierarchy in plant grain weight components can hence be interpreted as reflecting differences between active and passive plasticity. Moreover, because of the processes involved in responding to specific signals, active plasticity can come at a cost (DeWitt et al., 1998). Due to the sequential building of plant grain weight components in annual plants, it can further be hypothesized that the tradeoff between the cost of active plasticity and its advantages will be more beneficial for early processes than late ones, as illustrated by the tillering dynamics and the R:FR signaling. We also observed phenotypic plasticity in height during plant growth and it displayed a convergence pattern consistent with the shade avoidance syndrome, as already documented in barley varietal mixtures (Dahlin et al., 2020). Because this syndrome is associated with more biomass being allocated to vegetative structure to the detriment of reproductive ones, such a plasticity in height is considered harmful in crops managed with efficient herbicides (in that weed competition is not considered an issue), with ongoing efforts to reduce it in wheat (Wille et al., 2017). A reduced plasticity in stem elongation was for instance observed in modern, smaller cultivars in comparison to older, taller ones for oat (Semchenko and Zobel, 2005) and wheat (Weiss et al., 2009). Interestingly, we observed a similar convergence pattern in terms of tillering dynamics (Figure 4) as well as a significant relationship between the plasticity in tiller emission and height differential in mixture (Figure 7). This possibly reflects a causal interplay between height and tillering due to a shared mechanism, as exemplified by the pleiotropic effect of Rht genes (Xu et al., 2023). In terms of genericity, even though our results on tillering plasticity underlying the RYT variability were obtained on bread winter wheat, they are of interest to small-grains cereals in general. Moreover, considering active vs passive nature of trait plasticities should be helpful when designing mixtures and selecting cultivars to blend.

### 4.4. Significance for assembly rules and breeding strategies

Regarding assembly rules, farmers often avoid mixing cultivars of different height or earliness (Borg et al., 2018). However, our results show that the tallest cultivars at maturity or the earliest ones are not always dominating in the mixture. For instance, Kalahari had the highest final height in the height mixture but it did not dominate whatever the year. Similarly, Accroc and Aubusson were the earliest cultivars in the earliness mixture, especially the first year, but their dominance ranks this year were respectively second and third. Phenotypic data such as final height and heading date, easily recorded in pure stand trials and largely made available to farmers, may hence not be that informative to predict the outcome of competition in varietal mixtures, but this remains to be tested on a larger data set in terms of cultivars, mixtures and years. No definitive guideline for mixture assembly based on height and earliness can be drawn from our study, but it highlights that avoiding mixing cultivars of different height and earliness puts an unnecessary constraint on the set of possible mixtures to choose from.

Moreover, farmers in France base their choices of wheat cultivars on the results of field trials, notably with different management intensities (Loyce et al., 2012), yield being a major determinant and currently all trials only involve pure stands. As shown in Figure 2, yield differences between cultivars in pure stands could be informative of the magnitude of yield plasticity in mixtures, although the strength of the association depends on the year. The wealth of pure-stand yield data could hence be of interest to guide farmers in their assembly for mixtures. Given the potential benefits, this will have to be further tested on a larger data set.

Furthermore, our study highlights the importance of monitoring the plasticity of key traits such as tillering to decipher the behavior and performance of cultivars in mixtures. However, beyond cultivar-cultivar interactions, the observed patterns of plasticity in both tillering phases notably depended on the year (cultivar-by-cultivar-by-year interactions), which complicates the definition of mixture assembly rules. Hence, knowing the yield variability of cultivars in pure stands over various environments, although useful (Wuest et al., 2021), may not be sufficient. For instance, the plant grain weight in pure stands can be similar over years, while differing significantly in mixtures (eg. cultivers Belepi and Kalahari). Complex interactions between plants in mixture and climate make mixture performances hard to predict. The plant-plant interactions in mixtures (GxG) may strongly depend on abiotic conditions (GxGxE), leading to complex interactions hard to disentangle. This calls for systematically observing more distinct mixtures over several years and sites to fill such a gap of knowledge (Labarthe et al., 2021) Although it may be relevant to select against plasticity in height due to the shade avoidance syndrome (Wille et al., 2017), such a strategy may ultimately be counterproductive if plasticities in height and tillering have partially shared genetic architecture. (Barot et al., 2017) suggested that plasticity in some traits may be worth keeping: tillering may be one such example.

To breed for mixture yield, (Litrico and Violle, 2015) proposed to maximize the phenotypic variance in agronomic traits between mixture components, while minimizing it for interaction traits. They defined agronomic traits as the ones at least partially determining agronomic value (e.g., grain yield), and interaction traits as those involved in interactions with neighbors. As our study demonstrates, tillering fits both these definitions, hence questioning the applicability of this framework.

### 4.5. Perspectives and open questions

Because of the detailed monitoring of individual plants in mixed stands, it was only feasible to experiment with eight cultivars and two quaternary mixtures. Despite the small number of mixtures, the observed RYTs were representative of the variability in the literature (Borg et al., 2018). Binary mixtures are expected to ease the study of interactions, at the cost of restricting the number of observed cultivars. We made the choice to experiment with mixtures more complex than binary ones, to stick to the reality of farmers’ practices, but with more complex interactions to disentangle. Furthermore, similarly to other studies (Dahlin et al., 2020), we ignored the below-ground organs, even though root traits can contribute to a functional understanding of mixing effects (Montazeaud et al., 2018). However, adding this layer of phenotyping to our experimental design would have required the usage of a field-based, non-destructive root phenotyping protocol which remains to be developed. Our analysis of reaction norms for plant grain weight and how they are explained by phenotypic traits such as height and earliness is strongly based on ecophysiological knowledge. Beyond the active plasticity for tillering by the R:FR sensing, more and more other signals involved in plant-plant communication are described (Ninkovic et al., 2019; Subrahmaniam et al., 2018), and they certainly deserve more attention to understand the outcomes of plant interactions in mixtures.

Predicting mixture productivities will require a better knowledge of plant development in both pure and mixed stands, notably regarding the tillering process. Due to the phenotypic plasticity between stands, the cultivar-by-cultivar interactions, and their dependency on environmental conditions, observing mixtures will not be dispensable. From our results, a challenge would consist in phenotyping at high throughput the maximum number of tillers at stage BBCH30, possibly at several densities. In pure stands a UAV-based strategy seems feasible (Wu et al., 2022), but transposing it for a large number of mixtures in multi-environment trials will require the development of a precision sower, i.e. a programmable sower able to sow individual seeds from various cultivars according to a pre-specified layout, like what we have done by hand, if possible at a speed comparable to that of conventional sowers. Another strategy is to develop functional-structural plant models with an explicit tillering process (Blanc et al., 2021), allowing an *in-silico* exploration of mixture behavior after calibration on data as the ones we produced here. Combining both strategies, i.e., UAV-based phenotyping and process-based modeling, is already feasible for wheat in pure stands (Jégo et al., 2012; Yang et al., 2021), but remains to be explored in mixed stands.

## 5. Conclusion

Crop diversification is a critical leverage for the agroecological transition, and intra-specific diversification at the field scale, using cultivar mixtures, is increasingly used by farmers to stabilize yields and control diseases. However, there still is a knowledge gap on how to design efficient crop mixtures and further investigations are needed. We therefore developed an experimental and analytical framework to better study the plant-plant interactions underlying mixture performance. Based on eight wheat cultivars, we assembled two cultivar mixtures contrasted for height and earliness. This allowed us to decipher the importance of trait plasticity, and more specifically the early and active plasticity of tillering driving spike number which is a major yield component. Extending such an experimental study and coupling it to ecophysiological modeling and on-farm experiments will allow a better understanding of the complex genetic-by-environment interactions, and design resilient mixtures in a less predictable agro-climatic context.

## Acknowledgements

The authors thank D. Tropée and G. Vieceli for their valuable help and support with the field experiments, as well as T. Randrianarisoa, N. Vazeux-Blumental, H. Belcram, A. Postec, L. Malicet-Chebbah, B. Rouger, E. Forst, G. Van Frank, E. Blanc, A. Merbhène, M. Turbet-Delof, I. Goldringer, A. Hospital, M. Colas, F. Ammar, A. Sidik Meite, K. Ménard and N. Tinomme. The authors thank T. Moittie (Asur Plant Breeding), M. Balduz (Lemaire Deffontaines), C. Duquet and M. Prevost (Limagrain Europe), C. Michelet (RAGT2n) and S. Caiveau (Syngenta France SAS) for providing the seeds. The authors thank R. Rincent for providing the adjusted means of heading date and final height from the BreedWheat panel. The authors thank V. Allard and P. Roumet for fruitful discussions on topics related to this work, as well as the three anonymous reviewers for their particularly constructive feedback. This work was supported by the doctoral school FIRE of the Learning Planet Institute and the BAP department from INRAE (PerfoMix project).

## Supplementary Material

**Table S1.** Composition of both mixtures, as well as heading date (since January 1, DOY=day of the year) and height at maturity of the cultivars as obtained from a field trial from the BreedWheat project.

| Mixture | Cultivar | Maintainer<br>(registration year,<br>country) | Heading<br>(DOY) | Heading<br>(calendar<br>date) | Final height<br>(cm) | Awns |
| --- | --- | --- | --- | --- | --- | --- |
| **Earliness**<br>**mixture** | Accroc | RAGT<br>(2010, FR) | 128.4 | 5 May | 87.02 | Yes |
|  | Aubusson | Limagrain<br>(2001, FR) | 129.9 | 7 May | 87.03 | No |
|  | Bergamo | RAGT<br>(2012, FR) | 145.3 | 25 May | 92.17 | No |
|  | Expert | Syngenta<br>(2007, FR) | 144.1 | 24 May | 90.93 | No |
| **Height**<br>**mixture** | Bagou | Asur Plant Breeding<br>(2007, FR) | 136.7 | 16 May | 82.97 | No |
|  | Belepi | Lemaire Deffontaines<br>(2013, FR) | 138.1 | 18 May | 100.93 | No |
|  | Boregar | RAGT<br>(2008, FR) | 140.7 | 20 May | 84.08 | Yes |
|  | Kalahari | Limagrain<br>(2010, DE) | 141.9 | 21 May | 115.19 | No |

**Table S2.** Dates of sowing and harvest for plants grown in pure and mixed stands, as well as the dates of the stages ear 1cm (BBCH30) and heading (BBCH55) for plants grown in pure stands.

| Mixture | Cultivar | Sowing date |  | Ear 1cm date |  | Heading date |  | Harvest date |  |
| --- | --- | --- | --- | --- | --- | --- | --- | --- | --- |
|  |  | 2019 | 2020 | 2020 | 2021 | 2020 | 2021 | 2020 | 2021 |
| Earliness mixture | Accroc | 31 October | 5 November | 10 March | 13 April | 28 April | 25 May | 20 July | 02 August |
|  | Aubusson |  |  | 14 March | 09 April | 02 May | 31 May |  |  |
|  | Bergamo |  |  | 27 March | 13 April | 08 May | 31 May |  |  |
|  | Expert |  |  | 03 April | 17 April | 19 May | 31 May |  |  |
| Height mixture | Bagou |  |  | 17 March | 13 April | 19 May | 02 June |  |  |
|  | Belepi |  |  | 08 April | 20 April | 17 May | 02 June |  |  |
|  | Boregar |  |  | 11 April | 21 April | 18 May | 02 June |  |  |
|  | Kalahari |  |  | 28 March | 19 April | 17 May | 02 June |  |  |

**Table S3.**
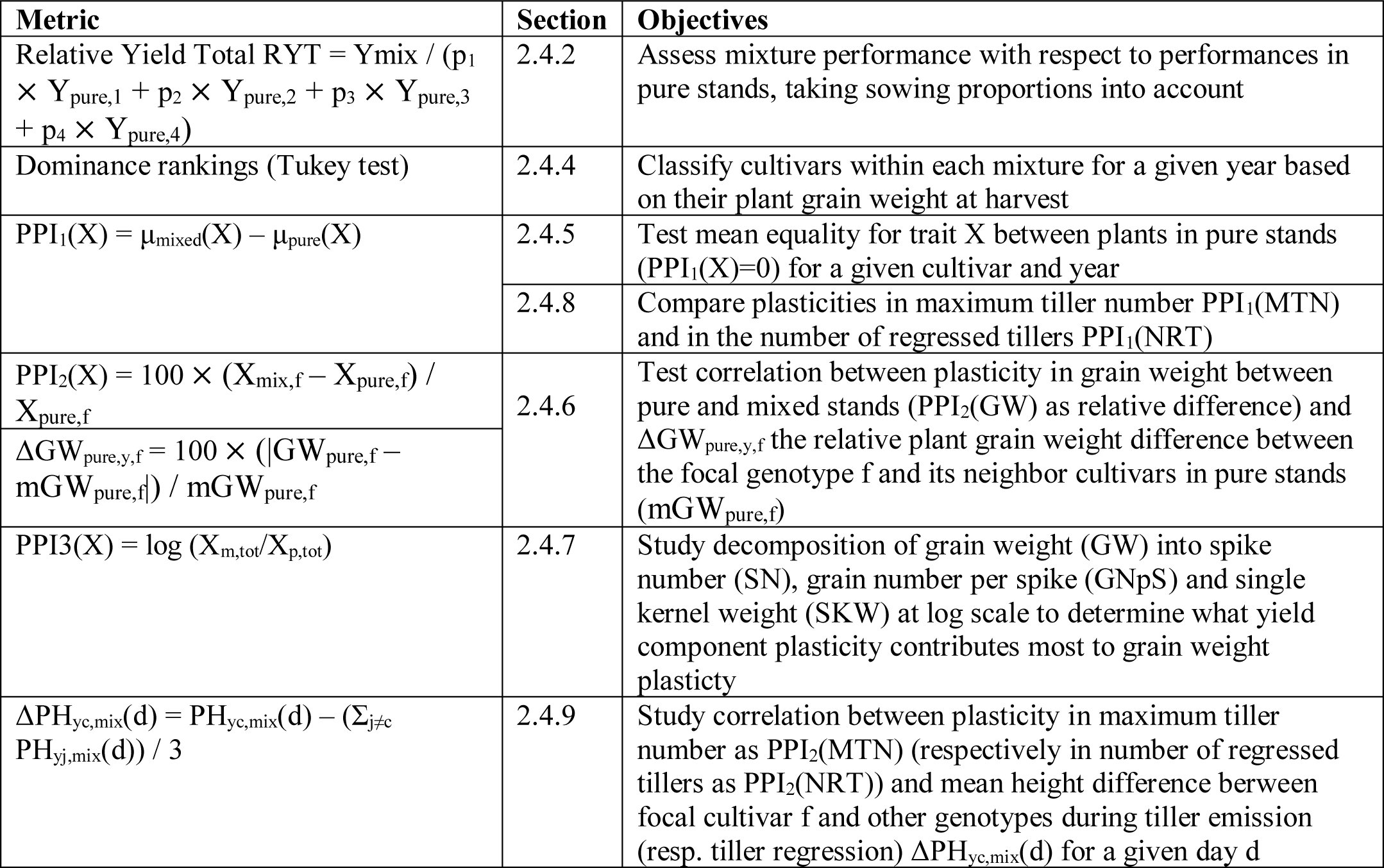
Summary of studied metrics and associated objectives

**Table S4.** Table with Tukey classes and matching dominance ranks for each cultivar over the two growing years.

| Mixture | Year | Cultivar | Letters from the Tukey tests |  | Dominance rank in mixed stands | Dominance status in mixed stands |
| --- | --- | --- | --- | --- | --- | --- |
|  |  |  | pure | mixed |  |  |
| Earliness mixture | 2019-2020 | Accroc | a | b | 2 | dominated |
|  |  | Aubusson | a | ab | 1.5 | dominating |
|  |  | Bergamo | a | b | 2 | dominated |
|  |  | Expert | a | a | 1 | dominating |
|  | 2020-2021 | Accroc | a | c | 3 | dominated |
|  |  | Aubusson | a | a | 1 | dominating |
|  |  | Bergamo | a | bc | 2.5 | dominated |
|  |  | Expert | a | ab | 1.5 | dominating |
| Height mixture | 2019-2020 | Bagou | a | b | 2 | dominating |
|  |  | Belepi | a | a | 1 | dominating |
|  |  | Boregar | a | c | 3 | dominated |
|  |  | Kalahari | a | bc | 2.5 | dominated |
|  | 2020-2021 | Bagou | b | a | 1 | dominating |
|  |  | Belepi | a | a | 1 | dominating |
|  |  | Boregar | b | b | 2 | dominated |
|  |  | Kalahari | ab | ab | 1.5 | ~dominated |

**Figure S1.**
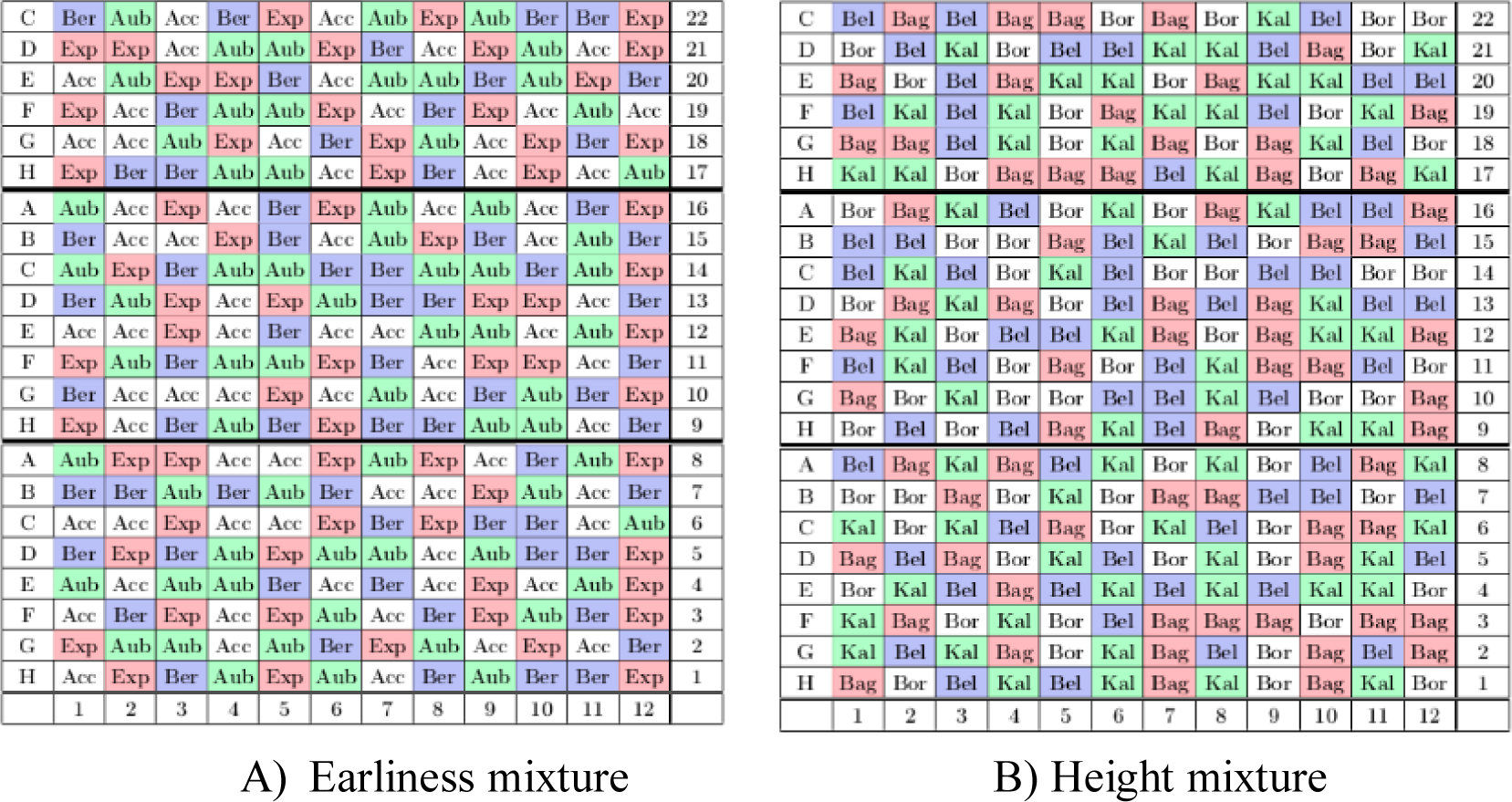
Unscaled schematic spatial distribution of cultivars in “nano-plots” for A) the earliness mixture and B) the height mixture for both years. Here, ranks are from 1 to 12 (x-axis) and rows are from 1 to 22 (y-axis). In each mixture there is one color per cultivar.

**Figure S2.**
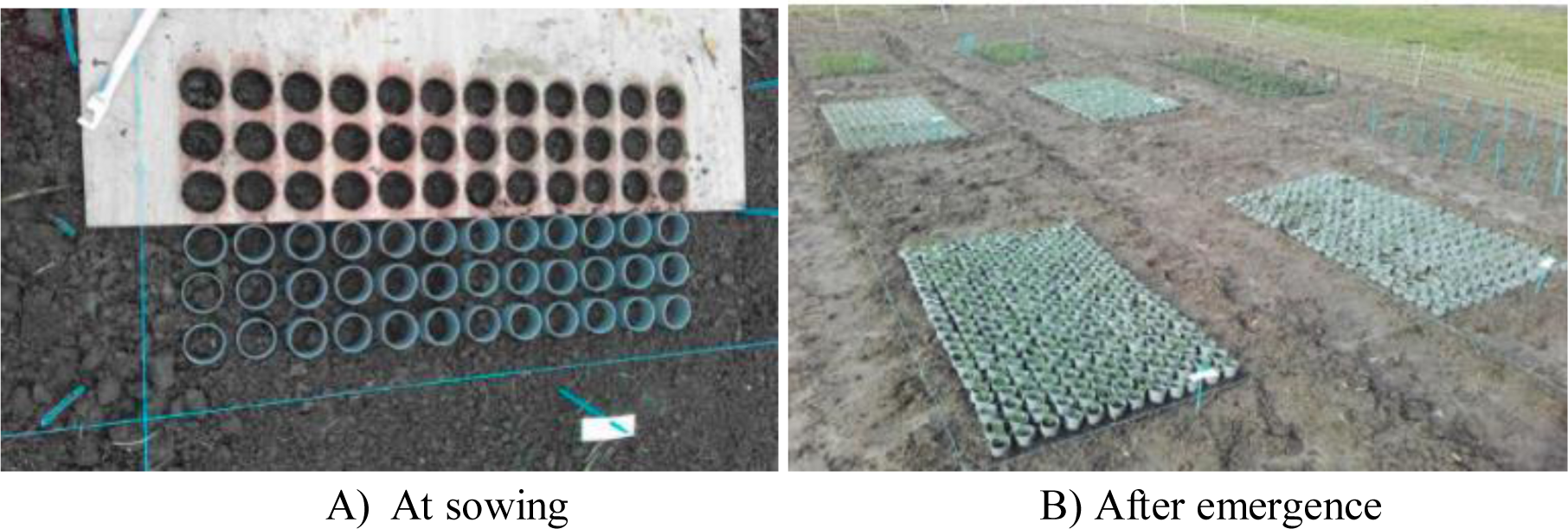
Images of “nano-plots” of mixtures with PVC rings in 2019-2020.

**Figure S3.**
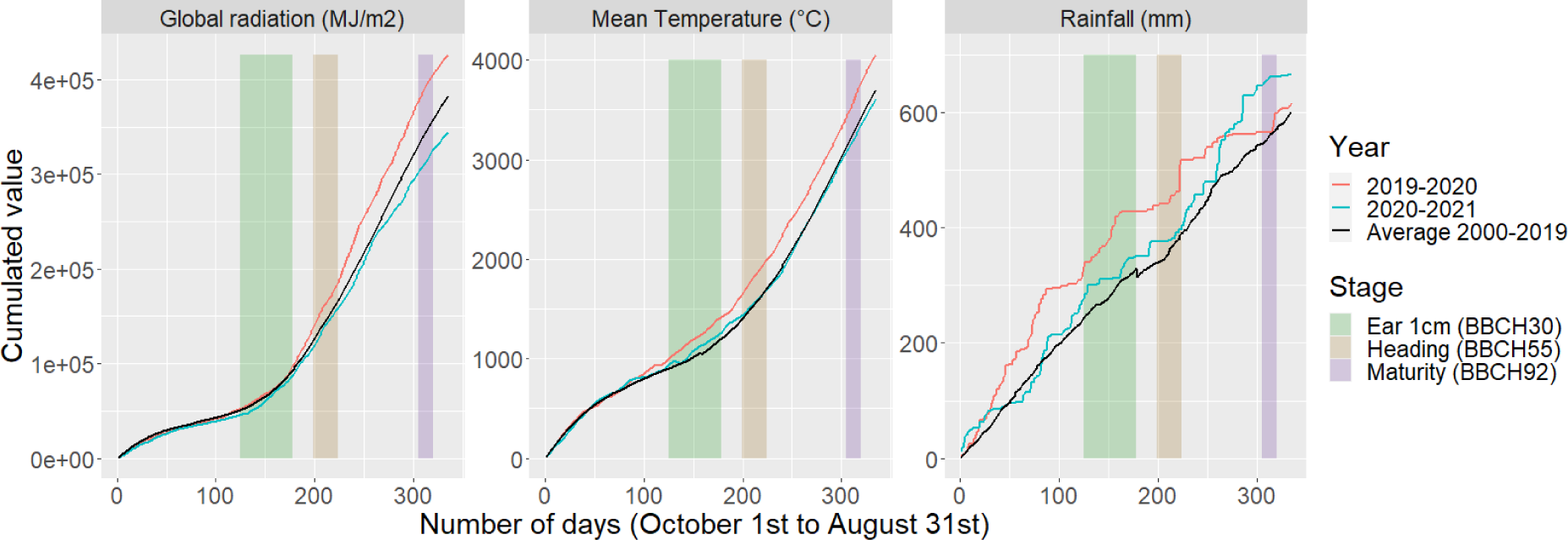
Cumulative daily global radiation, mean temperature and rainfall for both years, in comparison with their respective average from 2000 to 2019. Ribbons correspond to approximate ranges of the stages “ear 1cm”, “heading” and “maturity” which correspond respectively to BBCH30, BBCH55 and BBCH92.

**Figure S4.**
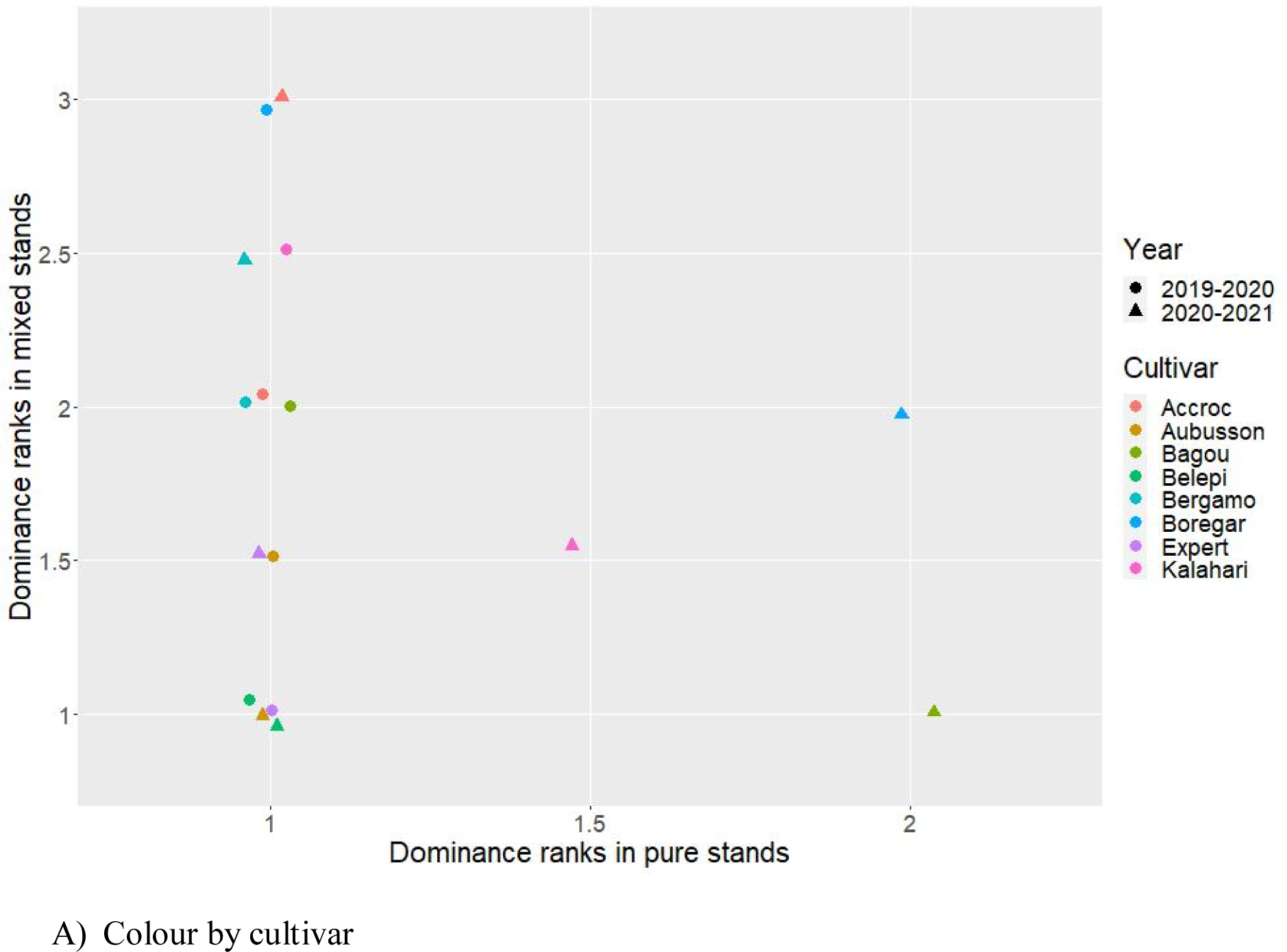
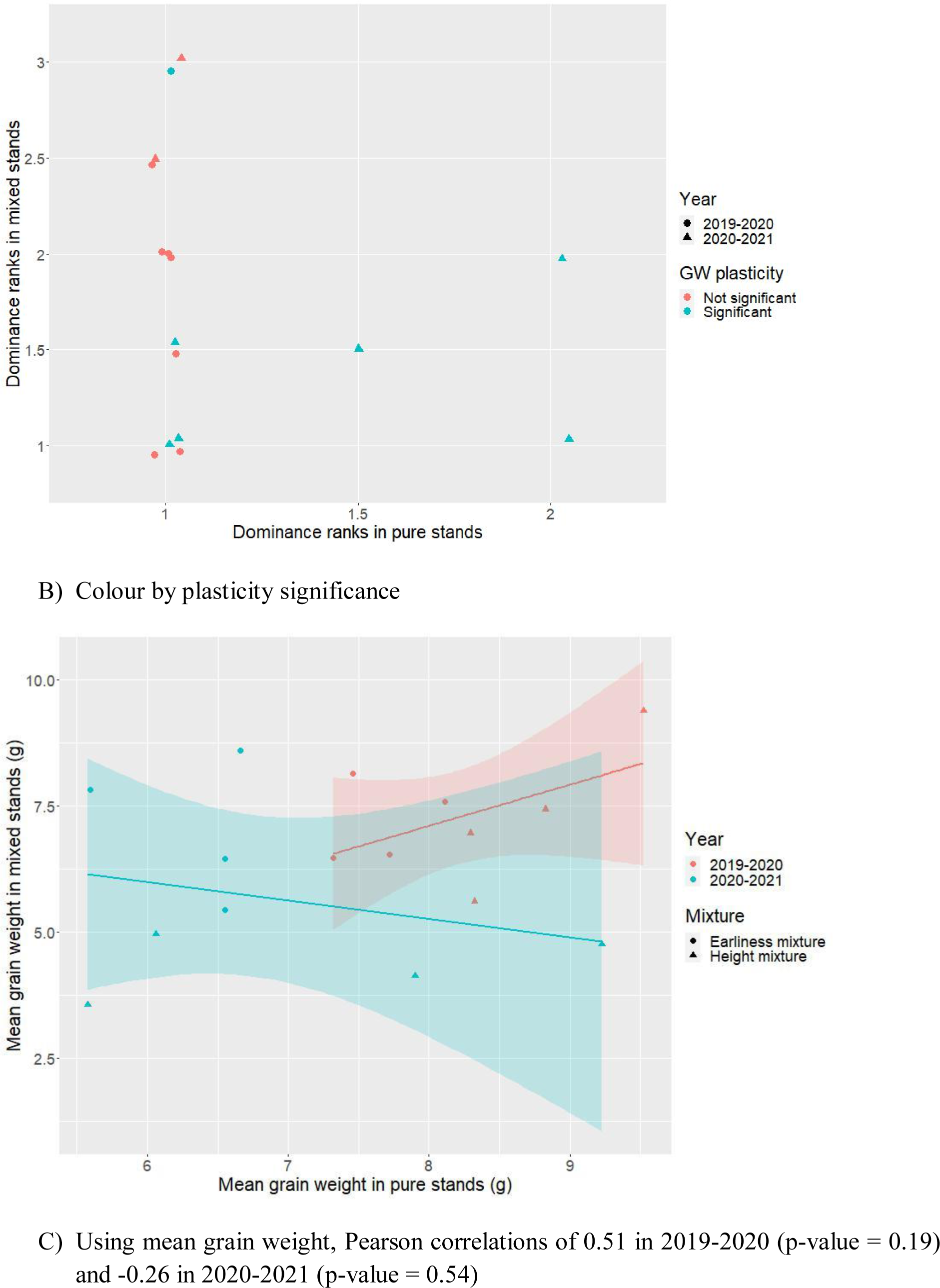
Dominance ranks (A,B) and mean grain weights (C) in pure vs mixed stands.

**Figure S5.**
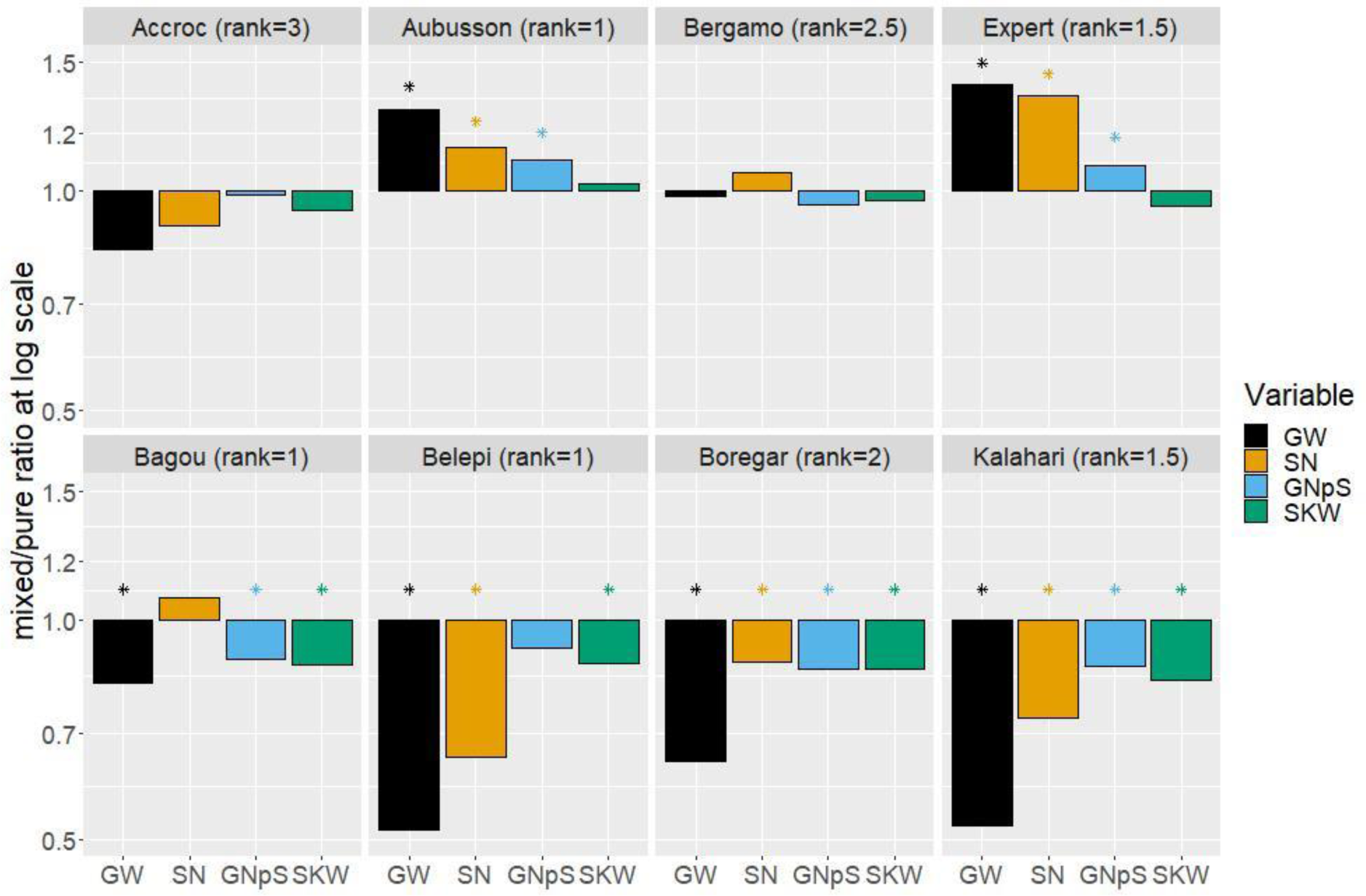
Log values of phenotypic plasticity of grain weight (GW) and its components, spike number (SN), grain number per spike (GNpS) and thousand kernel weight (TKW) in 2020- 2021. Stars indicate significant differences in pure versus mixed stands for a given variable, cultivar and year.

**Figure S6.**
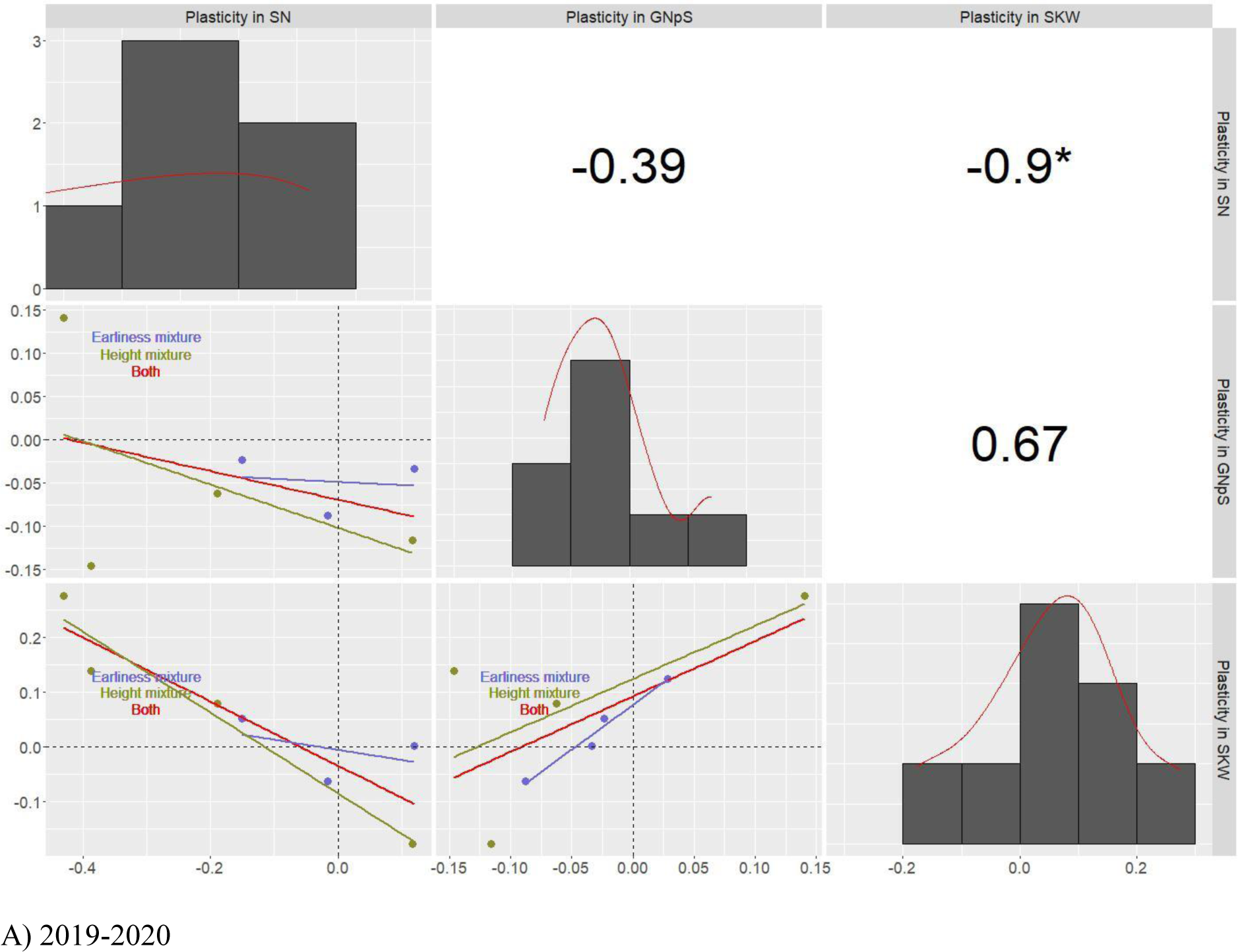
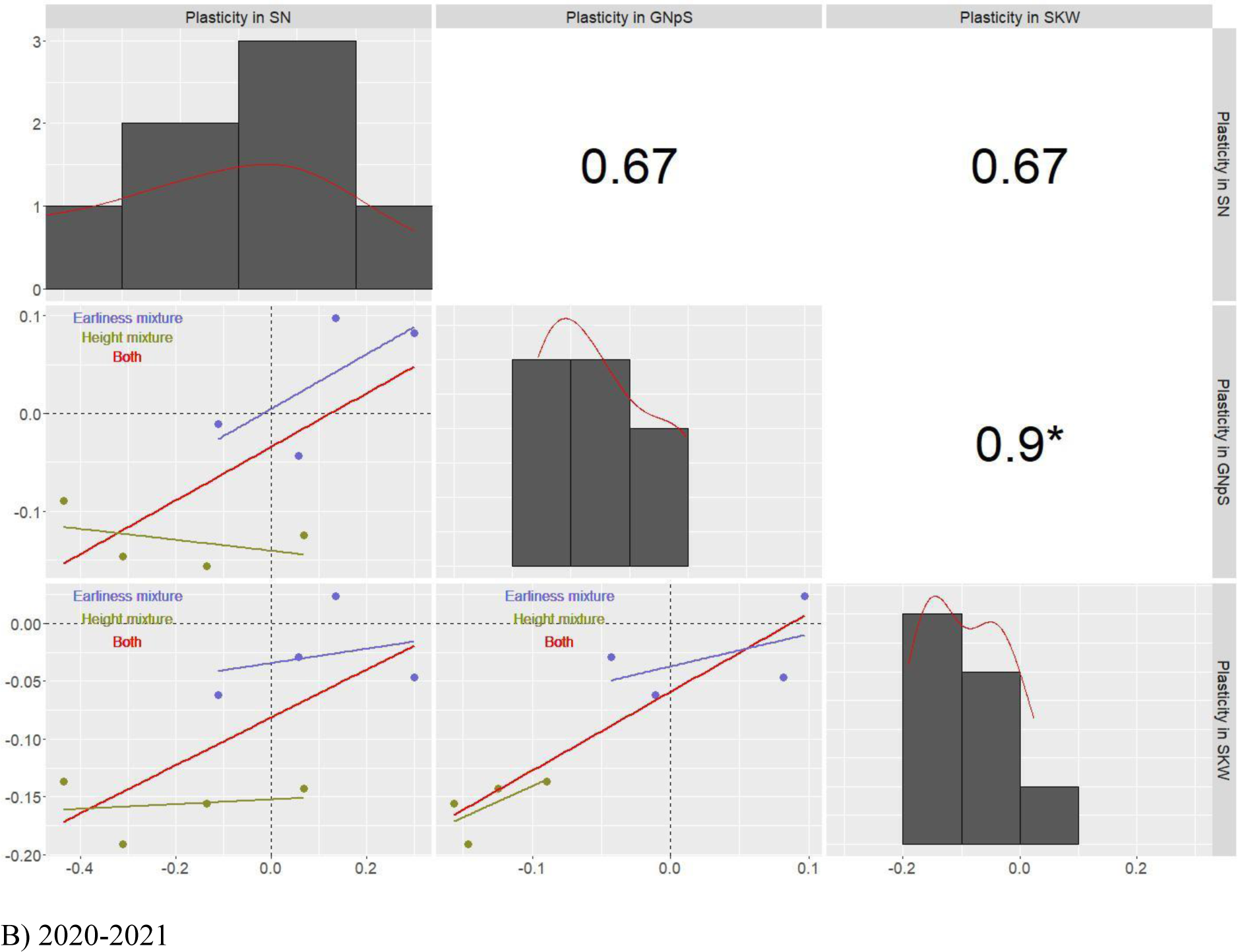
Distributions along the diagonal of the phenotypic plasticity in pure versus mixed stands for grain weight (GW) and its components, spike number (SN), grain number per spike (GNpS) and single kernel weight (SKW) in A) 2019-2020 and B) 2020-2021. Pairwise scatterplots are below the diagonal, in green for cultivars of the earliness mixture and purple for cultivars of the height mixture with corresponding linear regressions for each mixture and both (in red). Pearson correlations are above the diagonal, with stars to indicate significance at the 5% level.

**Figure S7.**
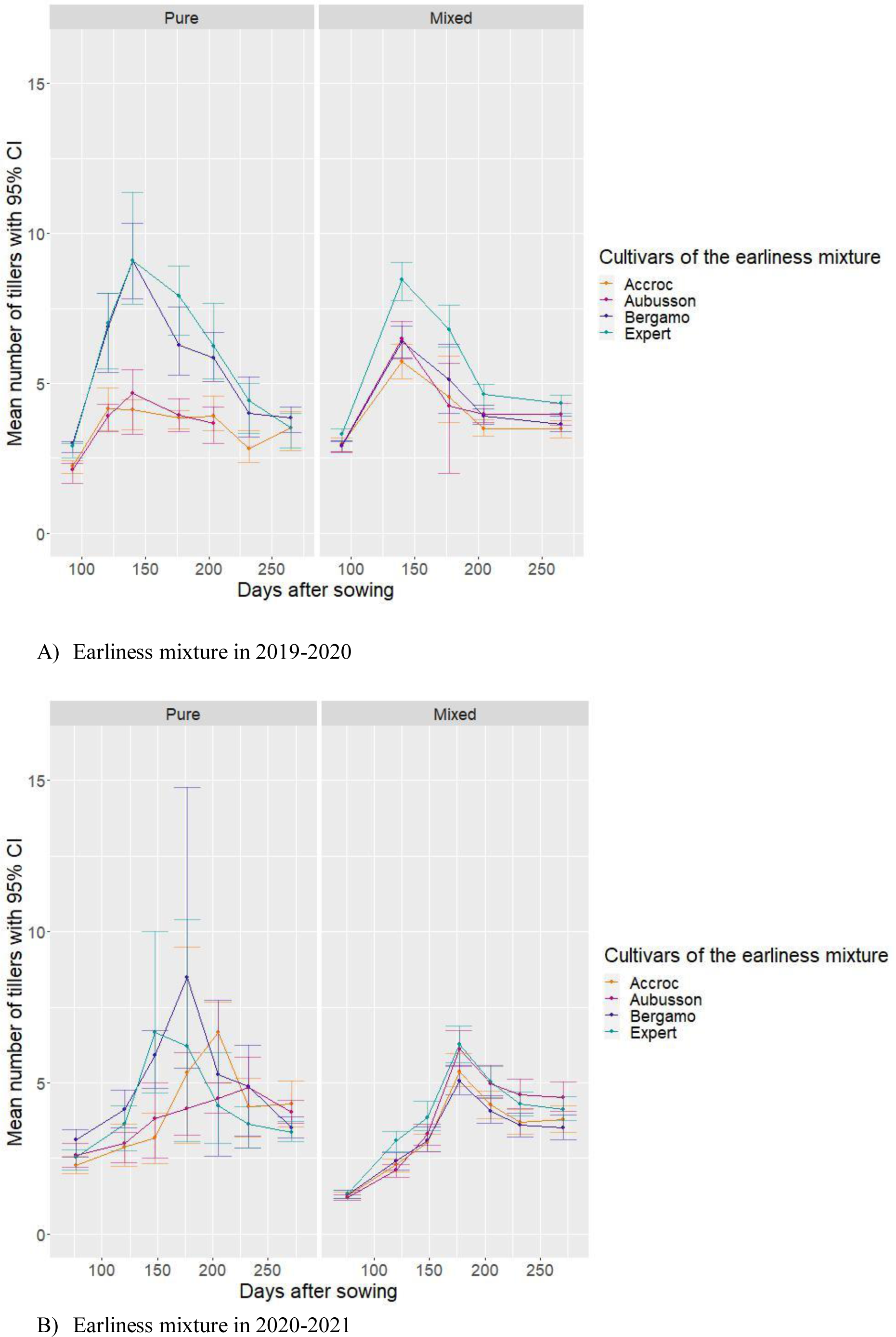
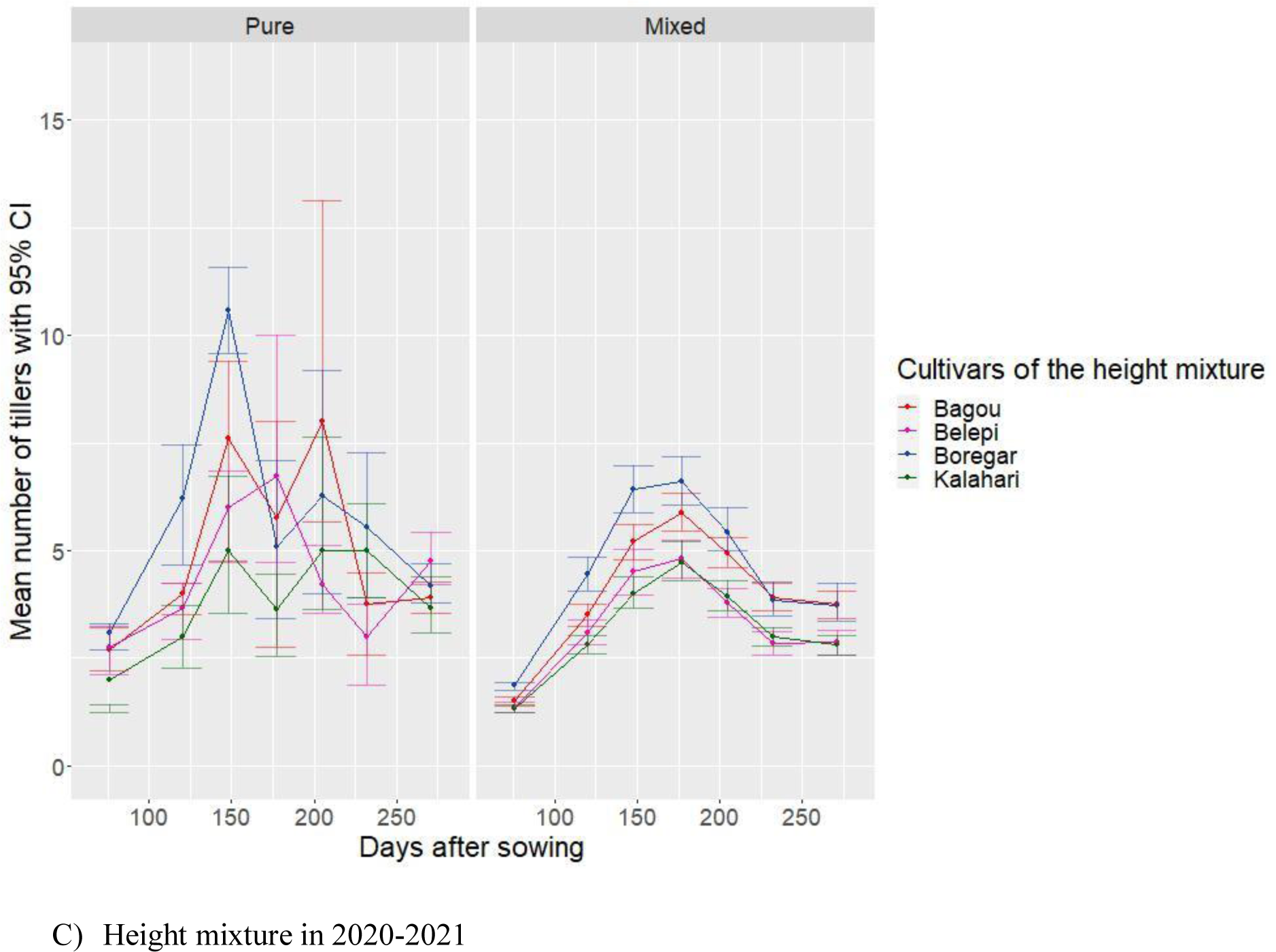
Average tillering dynamics from individual plant data all along the crop cycle for each cultivar of A) Earliness mixture in 2019-2020, B) Earliness mixture in 2020-2021, C) Height mixture in 2020-2021 for both pure and mixed stands. Error bars correspond to 95% confidence intervals.

**Figure S8.**
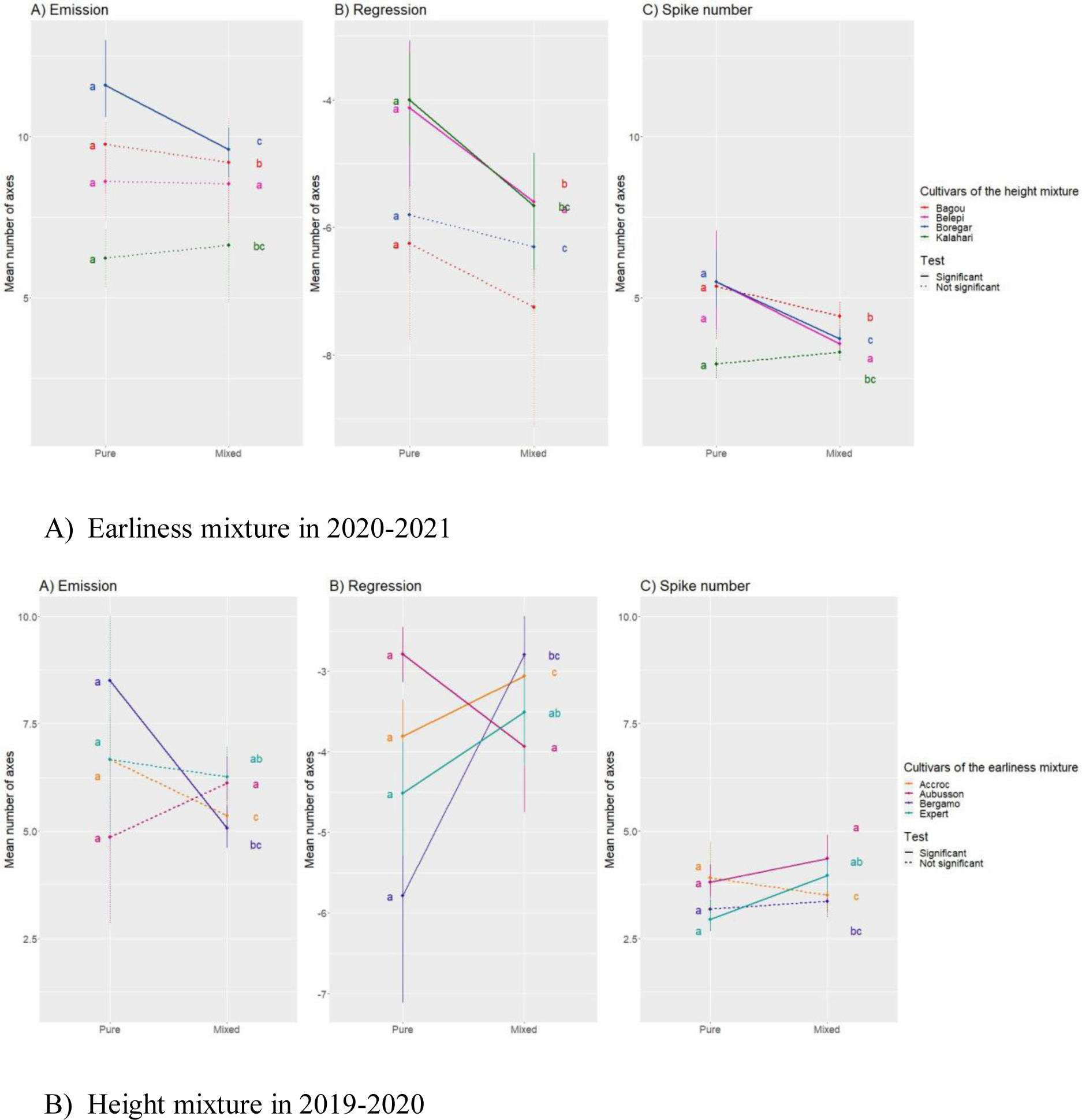
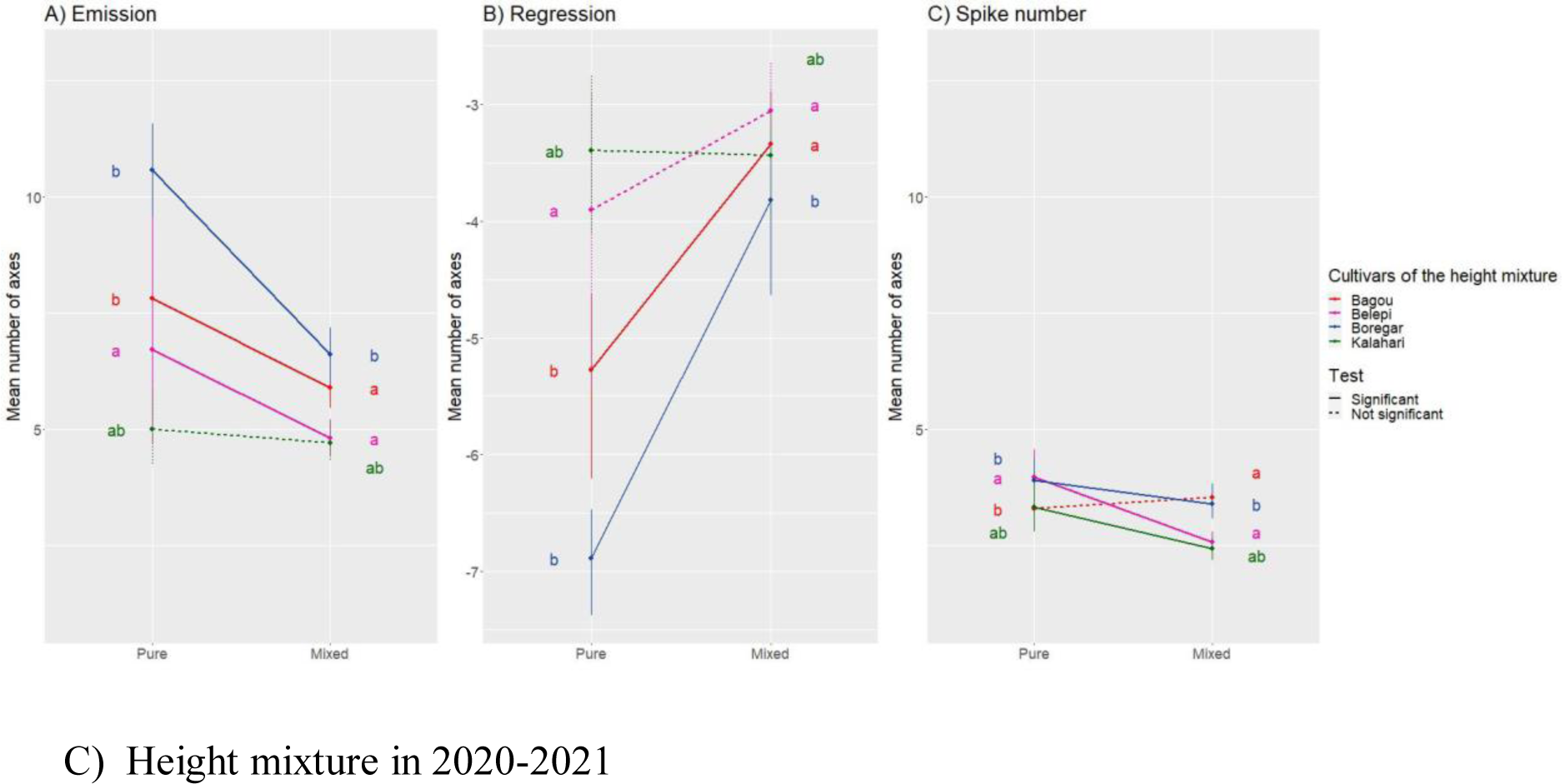
Reaction norms of the maximum tiller number (MTN) and the number of regressed tillers (NRT) during the emission and regression phases of axes per plant between pure and mixed stands per cultivar of the earliness mixture in 2019-2020 (A), as well as the height mixture in 2019-2020 (B) and 2020-2021 (C). Vertical segments represent confidence intervals at 95%. Letters indicate pairwise differences between cultivars in pure or mixed stands. Line types indicate the statistical significance of the reaction norms.

**Figure S9.**
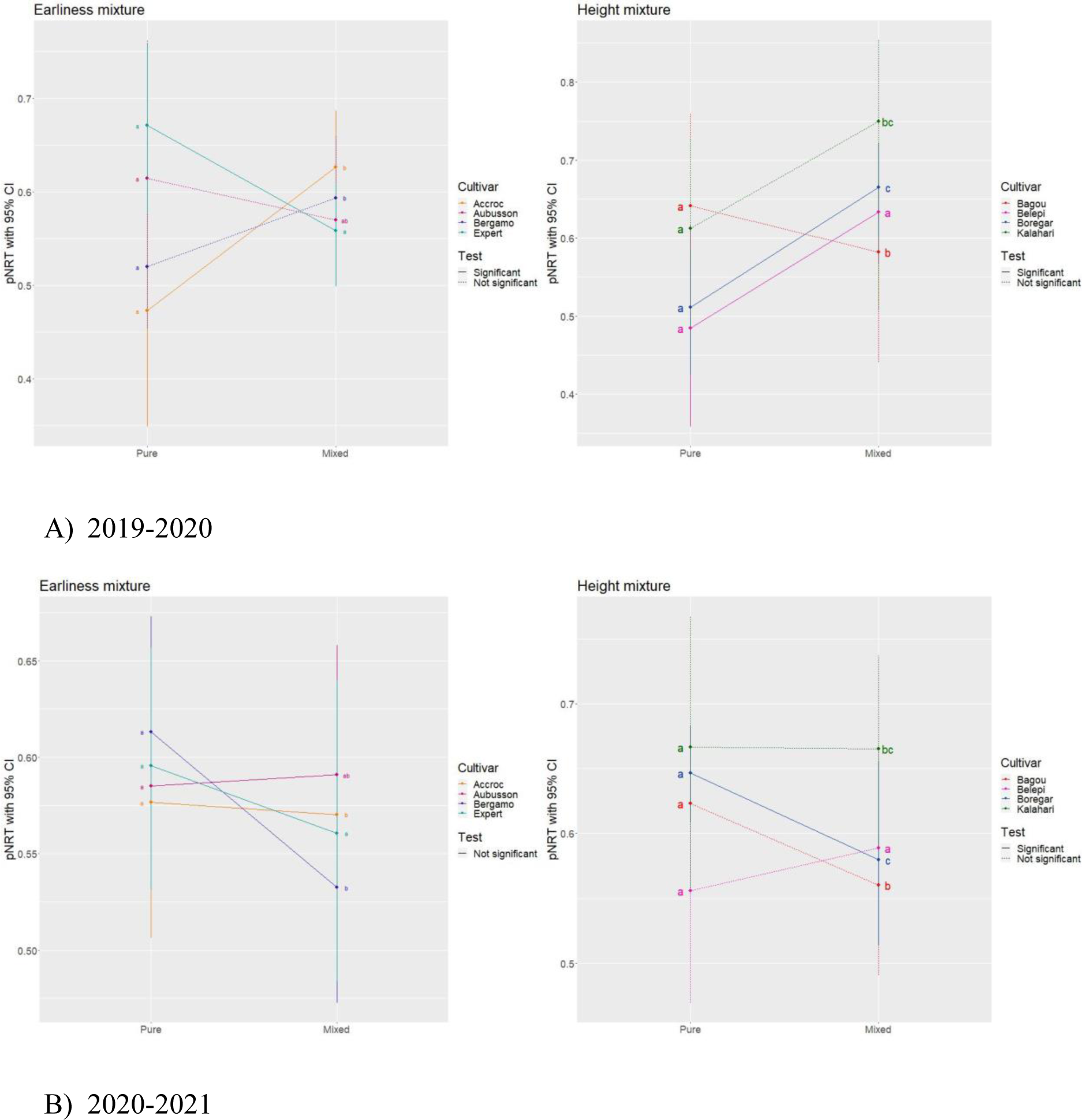
Reaction norms of the proportion of regressed tillers (pNRT) between pure and mixed stands per cultivar of the earliness and height mixtures in 2019-2020 (A) and 2020-2021 (B). Vertical segments represent confidence intervals at 95%. Letters indicate pairwise differences between cultivars in pure or mixed stands. Line types indicate the statistical significance of the reaction norms.

**Figure S10.**
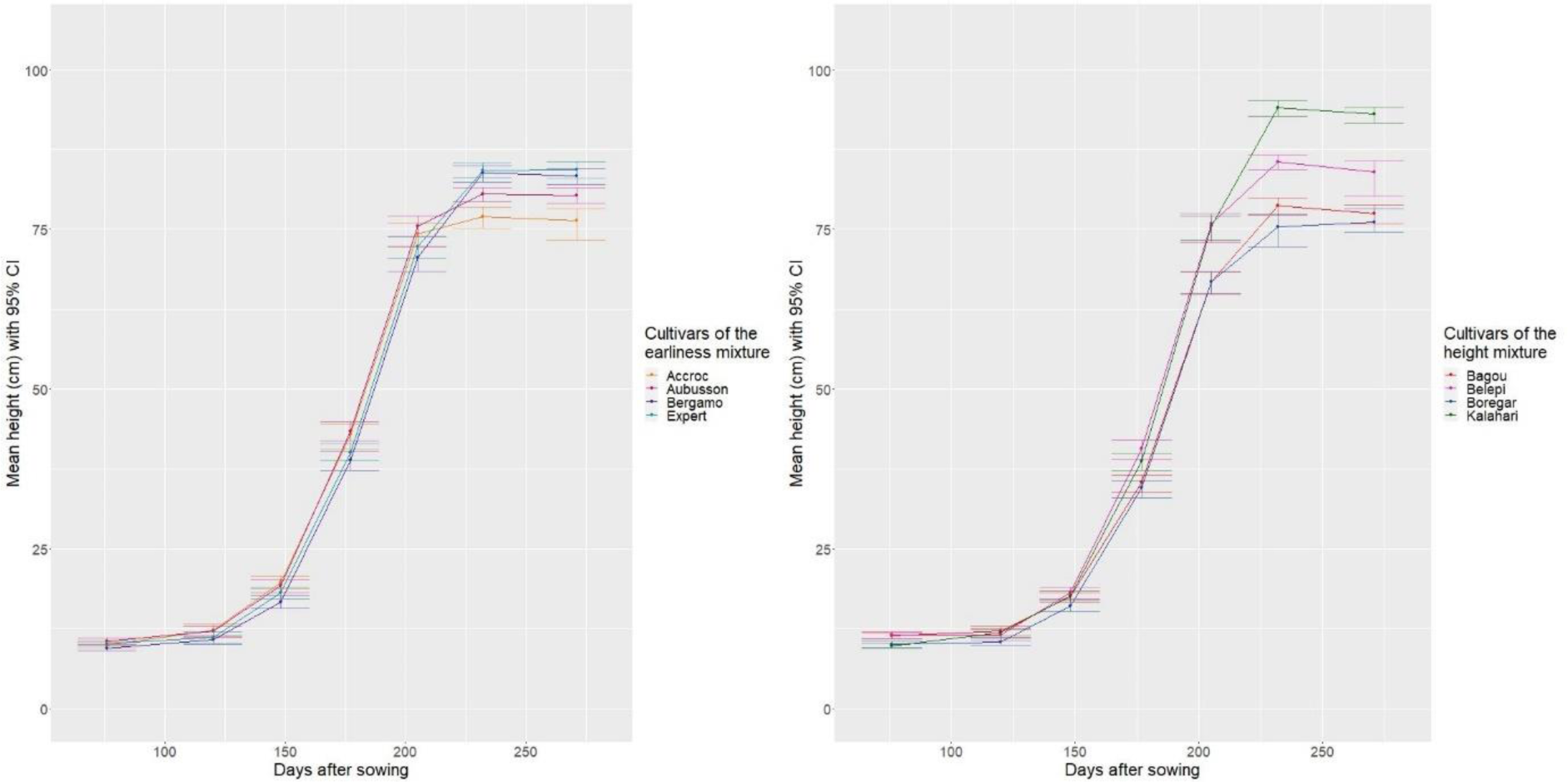
Plant height (in cm) throughout growth for the different cultivars grown in mixed stands for both mixtures in 2020-2021. Vertical segments represent confidence intervals at 95%.

**Figure S11.**
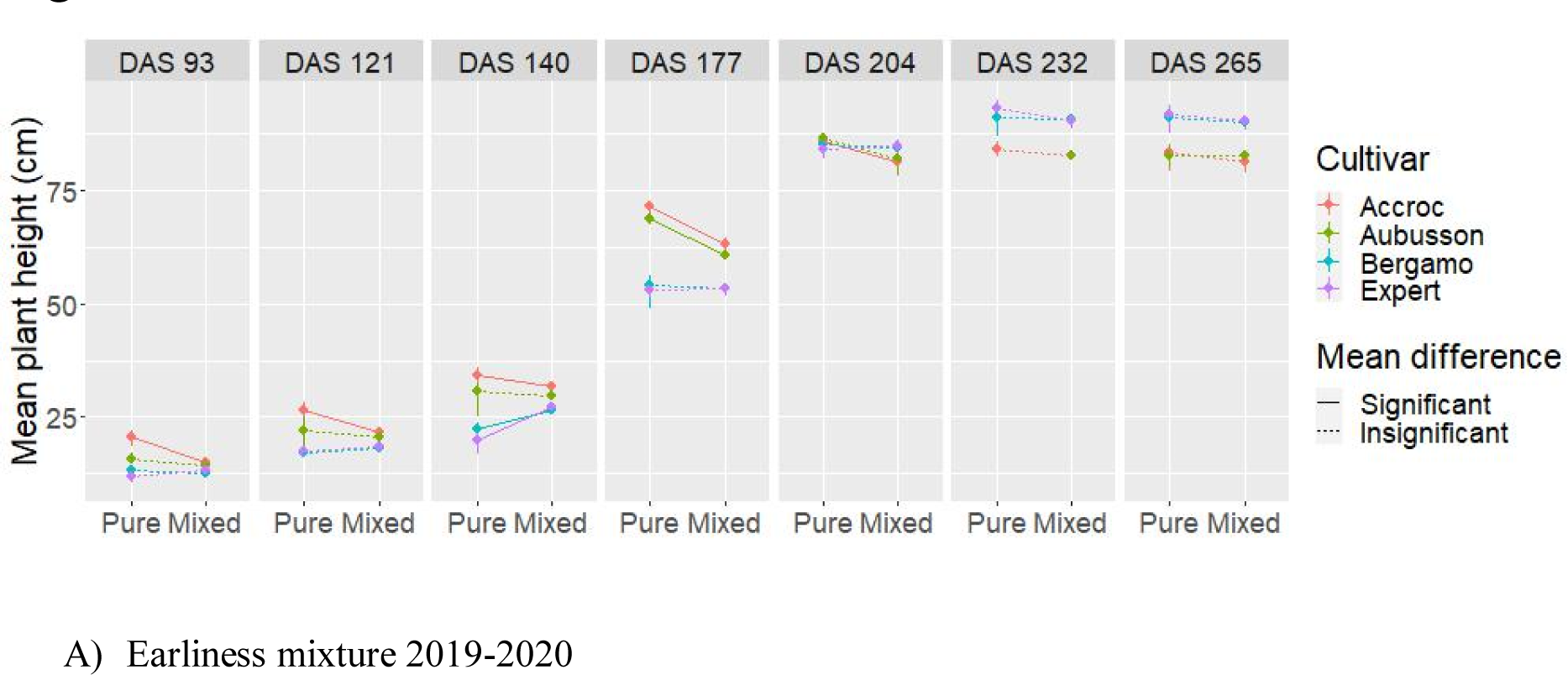
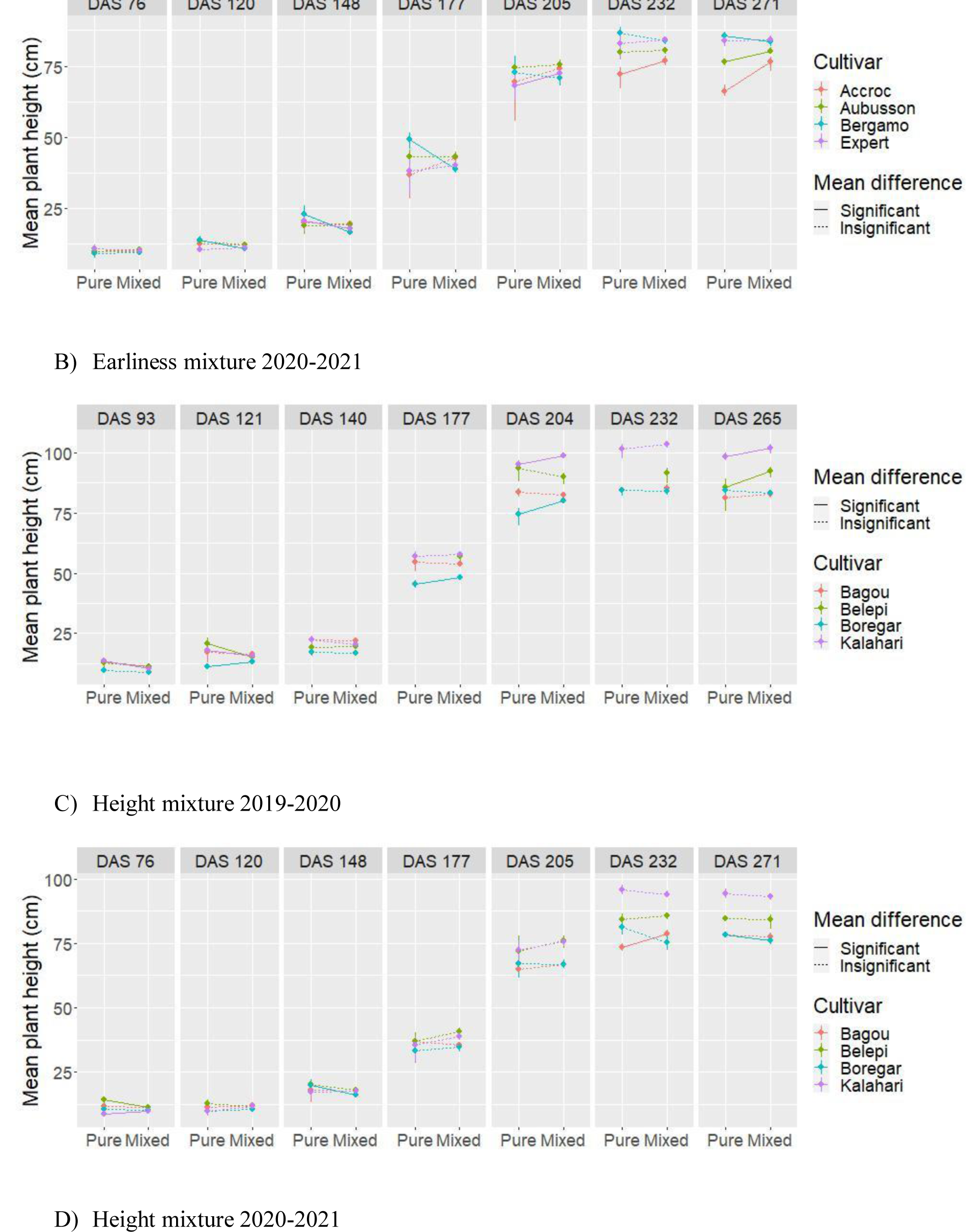
Reaction norms of plant height all along the crop cycle between pure and mixed stands per cultivar for both mixtures and years. Line types indicate the statistical significance of the phenotypic plasticities. “DAS” stands for “days after sowing”.

**Figure S12.**
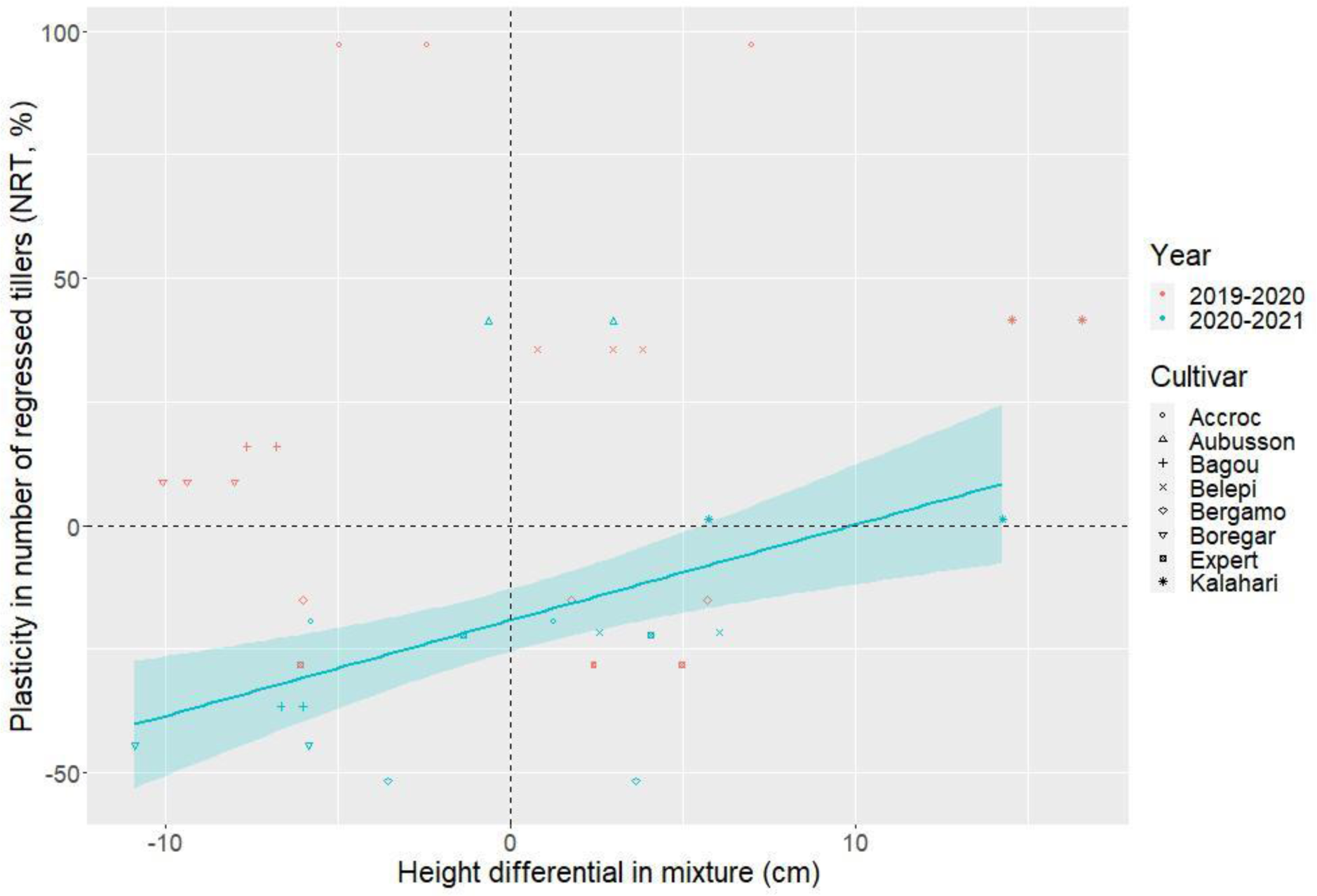
Plasticity of the number of regressed tillers (NRT) versus height differentials in mixture computed during the regression phase. Only significant regression lines are displayed. The height differential was significant at α=5% in 2020-2021: slope = -0.973 ± 0.627 in 2019- 2020 (p-value 0.126) and slope = 1.936 ± 0.516 in 2020-2021 (p-value 3.89 × 10^-4^, R^2^=0.18).

